# Human neutrophils release extracellular RNA that modulates *Aspergillus fumigatus* gene expression across kingdoms

**DOI:** 10.64898/2026.10.19.758605

**Authors:** Alexander Bruch, Lukas Schrettenbrunner, Bhawana Israni, Erik Böhm, Xiaoqing Pan, Corissa Visser, Thomas Orasch, Axel A. Brakhage, Matthew G. Blango

## Abstract

Extracellular RNAs (exRNAs) are emerging as potent bidirectional interkingdom effectors of host-pathogen interactions, particularly in plant and insect systems. Here, using small RNA-sequencing and genetic manipulation, we provide evidence consistent with a function of the human neutrophil exRNA-ome in modulating gene expression across kingdoms in the fungal pathogen *Aspergillus fumigatus*. The neutrophil extracellular space contains a diverse repertoire of RNA, which was corroborated using a droplet-digital stem-loop RT-PCR protocol to measure differential abundance of several tRNA-derived RNAs between two extracellular fractions. A closer inspection of the extracellular miRNA pool revealed enrichment of antimicrobial Let-7 family sRNAs and immunomodulatory miR-223, among others. Secretion of these sRNAs was largely impervious to infection status, suggesting that neutrophils produce and maintain an antimicrobial exRNA environment against *A. fumigatus* that requires further targeting. Cross-kingdom miRNA target prediction and exRNA delivery experiments revealed that putative fungal targets are modulated independently of the fungal argonaute proteins and in a donor-dependent manner. Although heterogeneous, for donors that exhibited a silencing effect, mutated target sequences in the fungus did reverse silencing. We hypothesize that the exRNA landscape produced by neutrophils contributes to their multifaceted role in antifungal defense.

**SIGNIFICANCE:** The human lung is regularly confronted with microbes, including several hundred asexual spores (conidia) of the mold *Aspergillus fumigatus* per day. These conidia are rapidly cleared in the healthy lung by phagocytes and other innate defenses; however, in immunosuppressed individuals, particularly those lacking neutrophils, *A. fumigatus* can germinate and establish invasive life-threatening infections. Here, we uncover a previously unappreciated mechanism contributing to neutrophil antifungal immunity, the transfer of Let-7 family microRNAs from human host to fungal pathogen to control pathogen gene expression. This first example of cross-kingdom RNA interference between human and mold bolsters the emerging concept that hosts and pathogens alike exchange RNA molecules as interkingdom effectors during the course of infection.

## INTRODUCTION

Interspecies communication relies on a variety of secreted molecules capable of traversing the extracellular space before uptake by recipient cells and modulation of gene expression. Remarkably, seemingly delicate RNA molecules were found decades ago to be enriched in the extracellular space (1, 2) and protected from RNase degradation either by (i) encapsulation in extracellular vesicles (EVs) (3–5), (ii) interaction with RNA-binding proteins (RBPs) (6, 7) or (iii) formation of RNase-resistant double-stranded RNA (dsRNA) structures (8, 9). Extracellular RNAs (exRNAs) consist of different non-coding/coding RNAs (e.g. rRNA and rRNA-derived RNAs [rDRs], tRNAs and tRNA-derived RNAs [tDRs], long non-coding RNAs [lncRNA], mRNAs, and microRNAs [miRNA]) and can serve as mediators of stress signalling or as effectors to alter gene expression of donor or recipient alike (4, 5, 10). These RNA-based intercellular communication mechanisms are surprisingly conserved and have been observed in numerous contexts (11–13).

Among the RNA species identified in the extracellular space, the miRNAs are the best studied. These small RNAs (sRNAs) produced in humans by the RNase III-type endoribonuclease DICER1 act as guides to direct the RNA silencing machinery containing an argonuate protein (e.g., AGO2 in humans) to appropriate, complementary target sequences (14). As extracellular cargo, miRNA are described as cross-kingdom RNA interference effectors (RNAi; reviewed in (15)). Although miRNAs are the best-studied extracellular cargo, rRNA-derived RNAs and tDRs like tRNA halves (tRH) and tRNA fragments (tRFs) are quite abundant, frequently dwarfing the miRNA population (16). The generation of tDRs relies on RNases like ANGIOGENIN, RNase T, and even DICER1 to recognize and cleave full-length tRNAs (17–22). Although the tDRs are detectable under non-stressed conditions, their biogenesis is often boosted during stress and disease (17). Similar to miRNAs, some tRNA-derived small RNAs like 3’-tRFs have been shown to interact with and guide AGO2 to silence specific target mRNAs (23, 24). tDRs also bear signalling potential in the immune response, *e.g.*, through an intricate role in T cell activation (25). These sRNAs can be secreted alone or in association with EVs and are of increasing diagnostic value for human disease (26).

Multiple studies have described extracellular RNA-dependent signalling between plants and fungal/oomycete pathogens, with defined mechanisms of transport and uptake for siRNAs/miRNAs. For example, the devastating plant fungal pathogen *Botrytis cinerea* encapsulates and traffics Bc-sRNAs in EVs to the plant host during infection relying on clathrin-mediated endocytosis to ultimately hijack the plant RNAi machinery to silence relevant immune genes (27, 28). Conversely, the plant generates and sends sRNA-carrying EVs that target specific virulence genes in the fungus via cross-kingdom RNAi (29, 30). Similar interactions were reported for insects and their fungal pathogens, e.g., *Anopheles stephensi* and *Beauveria bassiana*, where the host secretes highly conserved Let-7 miRNAs to interfere with fungal pathogenicity and the fungus traffics bba-milR1 to disrupt insect immunity (31, 32).

Observations of RNA-dependent interspecies communication between fungal pathogens and human cells are limited. For instance, work done on monocytes infected with *Candida albicans* revealed an interaction reliant on hsa-miRNA-24-3p (33). Toll-like receptor 4 (TLR4)-dependent recognition of the fungal pathogen by monocytes led to the secretion of the miRNA encapsulated in EVs, which were taken up by the pathogen and resulted in a knock-down of *SOL1*, a negative regulator of virulence-relevant fungal filamentation (33, 34). Many of these examples describe a substantial dependence of interspecies RNA-trafficking on extracellular vesicles, implying a mechanistic relevance on EVs to protect, transport, and target RNA molecules, especially during fungal-host interaction (35). EV-based cell communication also occurs between immune cells and fungal pathogens as was established for polymorphonuclear granulocytes (PMN; commonly referred to as neutrophils)-derived EVs with antifungal properties against *A. fumigatus* (36–38). These infection-induced EVs were able to prevent germination of the filamentous fungus in *ex vivo* treatment experiments. Although various antimicrobial proteins like Cathepsin G, Defensin 1, and Azurocidin were found by proteomics to be enriched in the antifungal EVs, the full repertoire of molecules (e.g., proteins, lipids, nucleic acids) contributing to this multifaceted effect remains incomplete (37).

Here, we isolated extracellular RNA from primary human neutrophils after *ex vivo* challenge with *A. fumigatus* conidia to define the exRNA-ome released in response to this WHO-critical priority pathogen. We observed a diverse extracellular RNA landscape, including miRNAs with known antifungal capacity. We then defined a representative case of cross-kingdom RNAi between the human let-7a-5p miRNA and putative *A. fumigatus* target *metE* using genetic manipulation of the target binding site in the fungus. These data suggest that neutrophils maintain an antimicrobial exRNA landscape against *A. fumigatus* that is targeted to the fungus upon recognition of infection.

## RESULTS

### Human polymorphonuclear leukocytes release heterogenous extracellular RNA

The RNA composition of the extracellular space is considered dynamic, adapting to environmental inputs with a tailored response of specific RNA species and isotypes (3, 39, 40). It is no surprise then that immune cells rely on such dynamism in defense against ever-changing microbes (25, 33). Here, we leveraged previous efforts in characterizing extracellular vesicles produced by human neutrophils against *A. fumigatus* (37) to define exRNA in two extracellular fractions during fungal infection (**Figure 1A**). We applied a differential centrifugation approach after infection that resulted in two sub-populations: (i) a pellet corresponding to molecules sedimenting at 19,600 x *g* (20k fraction) and (ii) the resulting supernatant (supernatant fraction). We sequenced the RNA cargo of fractions from uninfected and conidia-infected neutrophils from four healthy human donors. While the cellular RNA content displayed expected signals for both rRNAs and sRNAs, extracellular RNAs seemed to be highly enriched with 20-200 nt molecules, with more larger RNA species found in the 20k fraction (**Figure 1A**).

**Figure 1.**
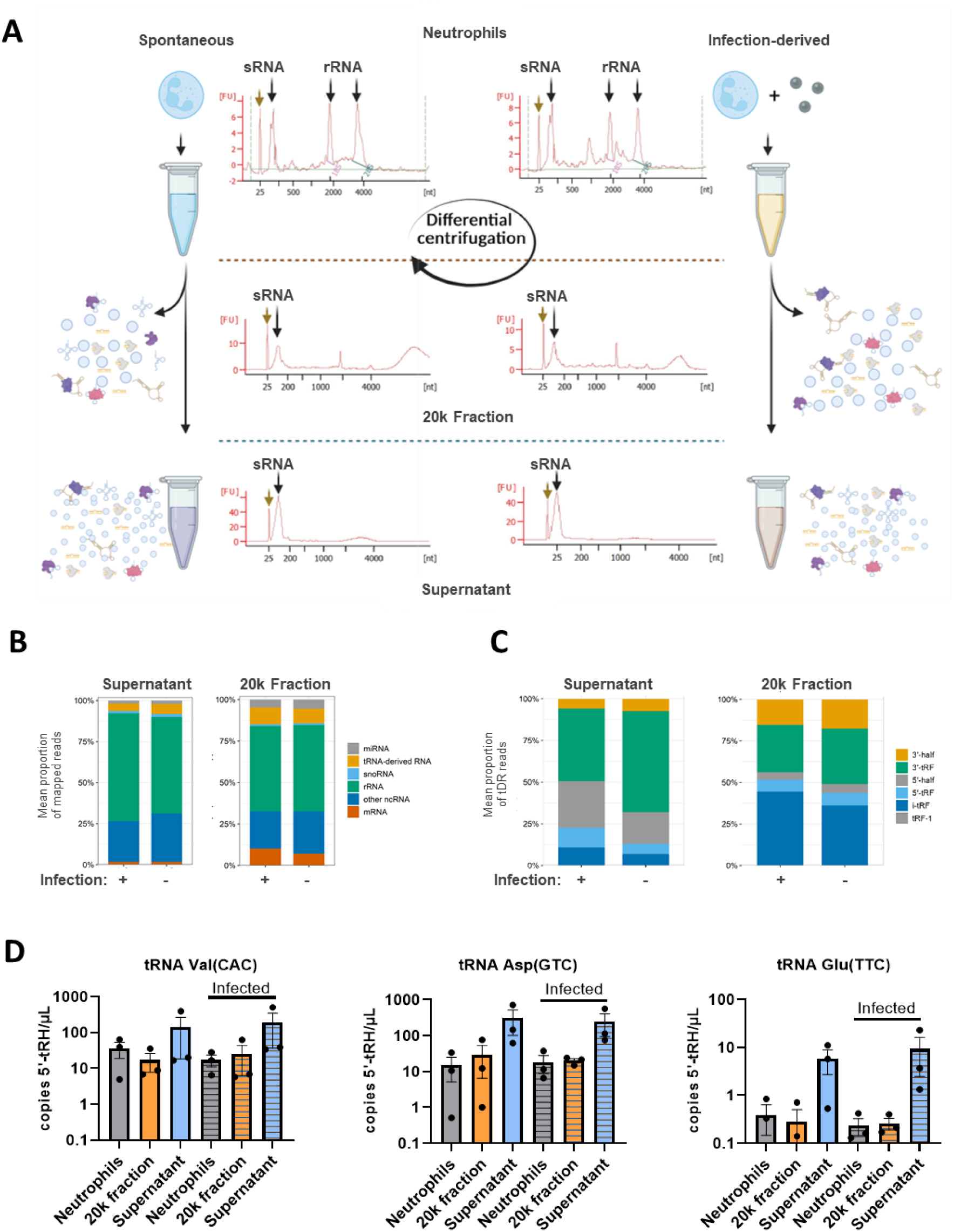
The diverse composition of neutrophil exRNA is enriched for miRNAs of the Let-7 family. (**A**) Model depicting the isolation modalities of extracellular RNA fractions in accordance to (37). RNA was isolated from pellets obtained at 19,600 x *g* centrifugation (20k fraction) and the corresponding concentrated supernatant (supernatant fraction). Bioanalyzer plots display RNA isolates from differently treated neutrophils and the respective fractions and conditions with rRNA and sRNA indicated. Created in BioRender, Blango, M. (2027) https://BioRender.com/uj92xgd. (**B**) Conducting small RNA sequencing with four biological replicates and subsequent *miRge3.0* analysis revealed the RNA species composition, and (**C**) tDR composition of the neutrophil extracellular space in uninfected and *A. fumigatus* infected conditions among both fractions. (**D**) Droplet-digital stem-loop RT-PCR of 5′tRNA halves (5-tRH) of tRNA Val(CAC) (*left*), tRNA Asp(GUC) (*center*), and tRNA Glu(UUC) (*right*). Striped bars indicate samples derived from infected neutrophils.

We performed small RNA-seq on the two extracellular fractions (as separate experiments described in methods) from infected or uninfected neutrophils using the Takara SMARTer smRNA-seq library preparation kit that uses template switching at the 5’ end of transcripts to minimize ligation bias. The resulting data was analyzed using the sRNA specific analysis software tool *miRge3.0* (41). Assessment with *miRge3.0* revealed diverse RNA species that differed minimally in biotype composition based on infection status (**Figure 1B, Table S1-2**) or donor (**Figure S1A-D**). The biggest groups were identified as rRNA- and ncRNA-derived transcripts, but we also observed stable fractions of miRNA and tDRs (**Figure 1B**). The largest differences were observed between the 20k and supernatant fractions, hinting at a potential difference between ectosome and exosome cargo, although the complex nature of the neutrophil extracellular environment during infection make this vesicle-focused distinction challenging to conclusively assess.

We observed a high relative proportion of lncRNA, particularly an abundance of Y RNA (e.g., Ro-Associated Y5) and rRNA molecules (e.g., 28S rRNA) in both fractions (**Figure S2A-B, Table S1-2**). The structural Y RNAs have previously been observed to be abundant in extracellular fractions (42–44), although their exact role in extracellular biology remains enigmatic. Using this approach, we observed some differential abundance of RNAs between the infection-derived and uninfected samples for the supernatant, but not 20k fraction, suggesting some discriminatory power in this fraction (**Figure S1C-D**). When we focused on only mRNA-derived transcripts in our extracellular fractions, we observed the presence of high levels of MT-RNR2, which encodes both the mitochondrial 16S rRNA and the small peptide, Humanin, particularly in the 20k fraction (**Figure S3A-B, Table S2**). The presence of MT-RNR2 was consistent with the literature highlighting a role for mitochondria in intercellular communication (reviewed in (45)). Beyond this mitochondrial rRNA, we observed some mRNA-derived transcripts that appeared to be slightly higher in abundance under infection derived conditions, e.g., *NBPF9-* and *ADARB2*-derived RNAs in the supernatant fraction (**Figure S3A, Table S1**), but more specific studies will be required to understand the function of such RNAs in extracellular biology.

### Extracellular tRF composition can be validated with droplet-digital, stem-loop RT-qPCR

Extracellular tDR patterns are known to reflect distinct cellular stresses and might therefore represent a potential readout for *A. fumigatus* conidia-infected neutrophils (16, 17, 21, 24, 25). To test this idea in our experimental system and provide corroborating validation of our sRNA-seq results, we performed a more in-depth analysis of tDRs under both tested conditions (**Figure 1C**, **Figure S4**, **Table S1-2**). The proportion of extracellular RNA corresponding to tDRs included both nuclear- and mitochondria-encoded molecules, with a specific enrichment of the latter in the 20k fraction (**Figure S4A-B**, **Table S1-2**). Interestingly, the detected tDRs displayed varying fragment species compositions (5′tRH, 5′tRF, 3′tRH, 3′tRF, tRF-1, internal-tRF [i-tRF]), with relatively more 5′-tRH in the supernatant fractions and i-tRF in the 20k fraction (**Figure 1C**, **Figure S4C-D**). In general, the composition and abundance of the different fragments varied only slightly depending on the tested condition (i.e. spontaneous [**-**] vs infection derived [**+**]) or between donors. Assessment of individual tRNA-derived RNAs from specific tRNA isodecoders allowed for discrimination of significant differences between infection-derived and supernatant fractions from untreated neutrophils (**Figure S4E-F**).

When we assessed the specific, abundant tDRs detectable in the extracellular fractions, we observed differences between the supernatant and 20k fractions (**Figure S5**). The most obvious difference was that the nuclear-encoded tDRs dominated the supernatant fraction, whereas mitochondria-encoded tDRs were most abundant in the 20k fraction (**Figure S5A,D**). The uninfected cell supernatant fraction were enriched for Arg-CCG 3′-tRF and SeC-UCA 3′-tRF, whereas slightly higher levels of Val-CAC 5‘-half were observed in the infected cell supernatants for the nuclear-encoded tRNAs (**Figure S5A**). A high abundance of Pro-UGG 3′-half was observed in both the infected and uninfected supernatant fractions for the mitochondria-encoded tRNAs, but the levels were elevated in the uninfected fraction (**Figure S5B**). In the 20k fraction, we saw the highest levels of Asp-GTC i-tRF and 3′-half from the nuclear-encoded tRNAs, whereas Leu-UAG, Gly-UCC, and Val-UAC i-tRFs dominated the mitochondria-encoded tRNAs, particularly in the infection-derived fraction (**Figure S5C-D**). The high abundance of valine tRNA fragments in both the supernatant and 20k fractions, particularly in the infection-derived supernatant fractions, led us to investigate this tRNA further. A closer inspection of the read coverage revealed that the valine tRNA halves are differentially enriched in the fractions, with 5’-tRH enriched in the supernatants and 3’tRF in 20k fraction, suggesting alternative trafficking upon cleavage (**Table S1-2**).

Continuing our investigation of extracellular tRFs, we utilized an elegant protocol reported in Tosar *et al*. (46) employing a stem-loop primer-based approach to reverse transcribe and quantify tRNA halves as validation of the dataset. Focusing on 5′tRHs, we tested several primers for RT-specificity and efficiency identifying halves derived from tRNA^Val^ , tRNA^Glu^ , and tRNA^Asp^ as promising candidates. We chose tRNA^Val^ due to its noticed differential fraction abundance, whereas tRNA^Glu^ , and tRNA^Asp^ were chosen for their lower abundance levels to test the sensitivity of our approach. Assessing equal amounts of RNA isolated from uninfected and *A. fumigatus-*infected primary neutrophils, as well as the respective 20k and supernatant fractions, indicated an enrichment for the respective 5′tRH in the extracellular fractions, particularly in supernatants (**Figure S6A-C**). We then performed absolute quantification of the tRFs with the help of droplet digital PCR (ddPCR) using the stem-loop RT-PCR strategy to define conditional differences of the intracellular and extracellular tRNA/tRF-ome and as a proof-of-principle detection of tRFs in the extracellular milleu, where we were able to confirm that the 5′tRHs of tRNA^Val^ , tRNA^Glu^ , and tRNA^Asp^ were enriched in the supernatant fractions (**Figure 1D**).

### Neutrophil extracellular RNA is enriched for Let-7 family miRNAs

We next focused on the extracellular miRNAs, as they are well-established in both antimicrobial defense and interkingdom communication (**Figure 1B**) (31, 33, 47). Evaluating the top twenty most abundant miRNAs revealed the 20k fraction to be enriched with miRNAs of the Let-7 family (e.g., let-7a-5p) (48), whereas the supernatant displayed a different picture with high levels of miR-223 and lower abundance of more diverse miRNAs like miR-23 and miR-92, among others (**Figure S7**, **Table S1-2**). No significant differences were detected in miRNA abundance between spontaneous and infection-derived exRNAs, which aligns with *in vitro*-studies of differentially stressed human cell lines displaying minimal alterations of extracellular miRNAs (39).

Studies in plants have shown specific RBPs like annexins (ANN1, ANN2) or argonaute (AGO1) to bind, enrich, and stabilize select extracellular miRNAs, suggesting RNA-protein interactions are a defining feature of exRNA compositions (30). Since primary neutrophils cannot be easily genetically modified to knock out/down human orthologues in our experimental set-up, we screened proteomics data from (37), and evaluated ANXA1, ANXA7, as well as ANXA11 for extracellular enrichment by western blot (**Figure S8**). All annexins displayed a remarkable extracellular enrichment under all conditions, although fungal infection boosted abundance, especially of ANXA1 (**Figure S8A**). Of note, the ANXA1 levels did exhibit some donor dependence (**Figure S8B**). In summary, the extracellular space of neutrophils contains a variety of small RNA species and RNA-binding proteins, and this composition remains largely stable throughout the tested conditions and between donors.

### Hyphal treatment with extracellular fractions results in decreased *metE* transcript

Small RNA-seq suggested a strong enrichment for miRNAs of the Let-7 family, especially in the 20k fraction (**Figure 1C**, **Table S2**). The 20k fraction was previously reported to bear multifactorial antifungal capacity by limiting the growth of *A. fumigatus*; however, the precise mechanism remains unclear (37). Studies of several different fungal infection models in plants and insects have reported exRNAs to be key factors to fight off fungal invasions by modulating virulence genes (29, 30, 32, 49, 50). This includes highly conserved Let-7 miRNAs conferring cross-kingdom function in the insect-fungal context and also parasitic worm and human antibacterial responses (32, 47, 51). We hypothesized that the increased Let-7 miRNA content in the 20k fraction could contribute to the antifungal capacity in a similar manner.

We began by testing the *sec2* (*AFUB_042620*) gene of *A. fumigatus*, since its ortholog in *B. bassiana* was previously shown to be silenced by a mosquito Let-7 miRNA by cross-kingdom RNAi (32). 22-h-old *A. fumigatus* hyphae (wild type, CEA10) were treated for 2 h with either 1X PBS (mock) or with freshly isolated 20k or supernatant fractions of the spontaneous or infection-driven condition, followed RT-qPCR for the targets. Relative transcript quantification resulted in significant silencing of *sec2* by the infection-derived 20k fraction (**Figure 2B**). Although not yet fully characterized in the *A. fumigatus* CEA10 lineage, deletion of *sec2* in strain IFM40808 resulted in several growth defects, including on rich media (52). We then deleted *sec2* in CEA17Δ*akuB^KU80^*, which resulted in decreased growth on malt agar, under 1 mM diamide-induced oxidative stress, on 0.5 µg/ml caspofungin-containing AMM, as well as a combination treatment thereof (**Figure S9A-B**), confirming an important role for Sec2 in the filamentous fungus and potential consequences as a cross-kingdom RNAi target.

**Figure 2.**
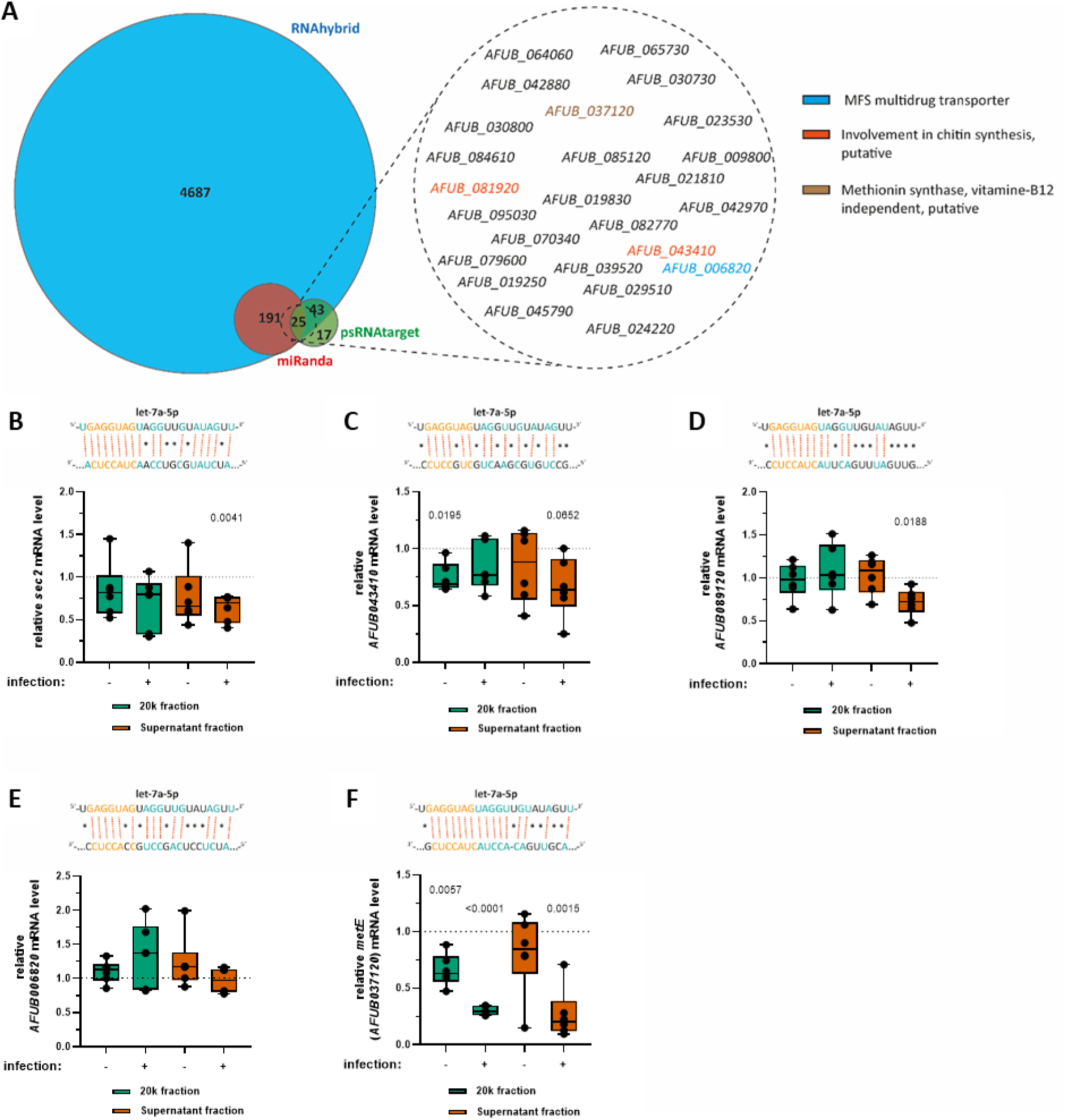
*metE* transcript levels are strongly decreased upon treatment with neutrophil extracellular fractions. (**A**) The Venn diagram displays the results of the miRNA target prediction tools RNAhybrid (e = −20, gap = 2-8; (53)), miRanda (S = 155, E = −20; (54)) and psRNAtarget (default parameters of Schema V2, 2017 release; (55–57)) used with the identified human miRNAs against the *A. fumigatus* A1163 (ASM15014v1) cDNA library. The bold number indicates the overlapping predictions. The Venn diagram was generated with (89). (**B-F**) Wild type (CEA10) hyphae were treated after 22 h growth with either 1X PBS (mock; dotted line), freshly isolated 20k fractions, or supernatant fractions from uninfected (**-**) or infected neutrophils (**+**) for an additional 2 h. The treated mycelium was washed, harvested, and subjected to RNA isolation, cDNA synthesis, and qPCR to quantify *sec2*, *AFUB_006820*, *AFUB_043410*, *AFUB_081920*, and *AFUB_037120* (*metE*). The measured values for all five genes were normalized to *cox5* transcript levels (88). Statistical significance was calculated using a one-way ANOVA test with a subsequent Dunnett’s posttest to control for multiple comparisons. A *p*-value ≤ 0.05 was considered significant.

Combining three different sRNA-target prediction tools, RNAhybrid (53), miRanda (54), and psRNAtarget (55–57), we predicted let-7a-5p miRNA binding sites in *A. fumigatus* cDNA, (**Figure 2A, Table S3**). While the RNAhybrid approach resulted in an uninformative 4,946 potential targets even after strict filtering, miRanda and psRNAtarget displayed fewer hits (216 and 85, respectively), with an overlap of 25 common predictions. Interrogation of the overlap revealed genes responsible for a wide range of functions in the fungus with no particular enrichment (**Table S3**). We chose four potential targets, encoding cell wall biosynthesis enzymes (*AFUB_081920*, *AFUB_043410*), a putative cobalamin-independent methionine synthase (*AFUB_037120*, renamed *metE*), or a multidrug transporter of the major-facilitator-superfamily (MFS, *AFUB_006820*) for RT-qPCR. Using the same setup as for *sec2*, we measured gene expression after delivery of extracellular fractions and observed similar slight transcript decreases (by ∼20%) after treatment with the spontaneous 20k fraction (*AFUB_043410*, **Figure 2C**) or the infection-derived supernatant fraction (*AFUB_081920*, **Figure 2D**). The mRNA levels of *AFUB_006820* displayed no significant change upon any treatment (**Figure 2E**). Interestingly, quantification of the *metE* mRNA levels indicated the strongest detected alteration (**Figure 2F**). Hyphae incubated with the 20k fraction from both conditions and the infection-derived supernatant fraction displayed strong *metE* mRNA reductions, which was especially pronounced with samples isolated from infected neutrophils (∼70-80% less transcript level compared to the control). Although *metE* is not the major methionine synthase gene of *A. fumigatus*, we suspected that its alteration might show some phenotypes under specific conditions. Upon creating a knockout, we revealed that gene deletion led to susceptibility when grown with a combination of stressors, for example with oxidative (e.g., 1 mM diamide) and cell wall stress (e.g. 0.5 µg/ml caspofungin) as well as with caspofungin treatment alone (**Figure S10**).

### Inactivation of the fungal RNAi machinery does not restore the *metE* mRNA level

The substantial decrease of *metE* transcripts after treatment with both infection-derived fractions may hint to an involvement of a cross-kingdom RNAi mechanism. Inspired by the literature, we rationalized two potential options on how Let-7 miRNAs might be trafficked and taken up to influence the fungal transcriptome (**Figure 3A**) (28, 30, 32, 58, 59). The first scenario is that extracellular let-7a-5p (and other miRNAs) are bound by human argonaute proteins to form functional ribonucleoprotein complexes (RNPs) that are taken up by the fungus to silence specific target genes like *metE* (**Figure 3A, I.**). This mechanism would rely only on trafficked host components and function independently of the fungal RNAi machinery.

**Figure 3.**
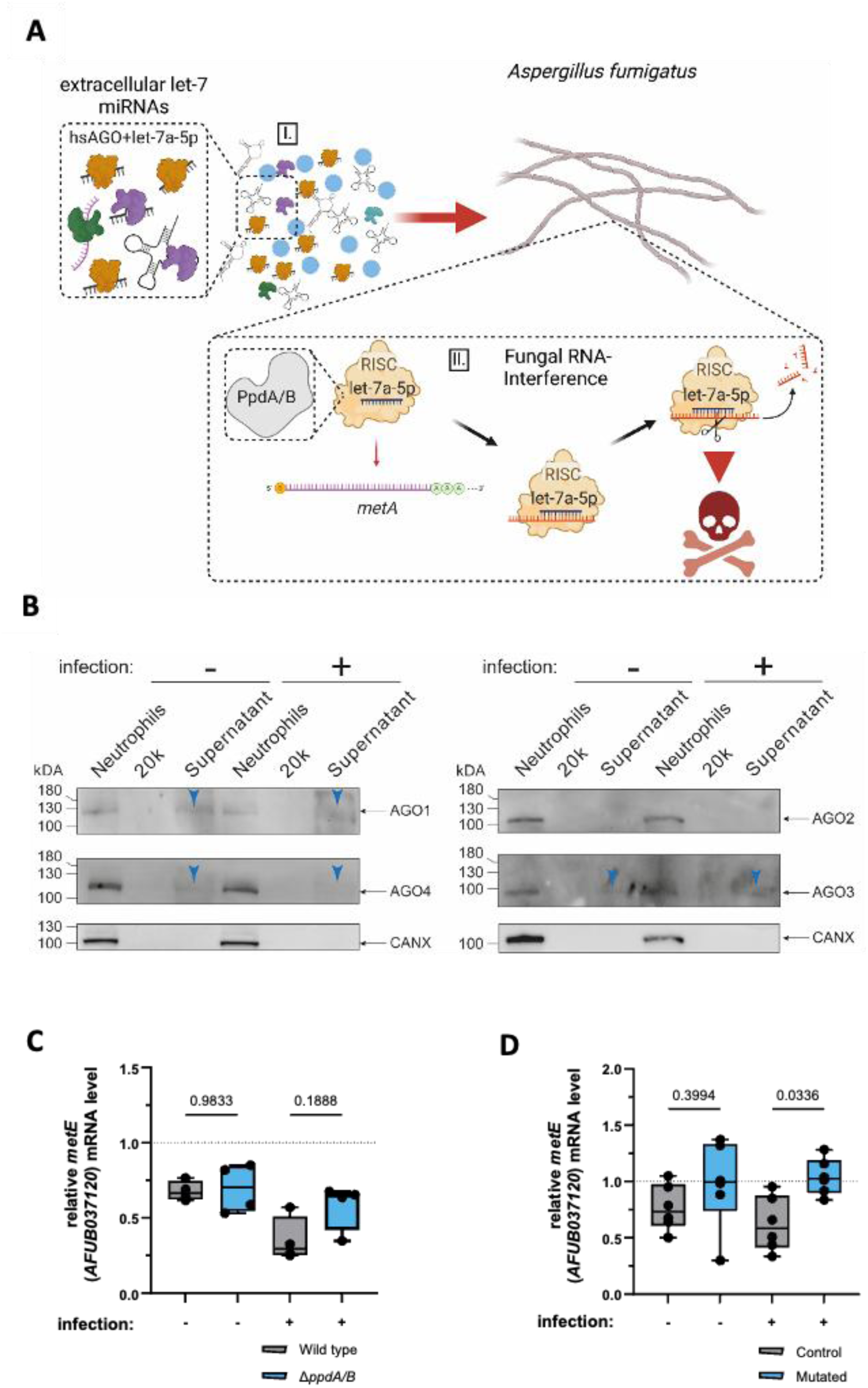
Mutation of target site seed sequence in *metE* transcript alleviates silencing. (**A**) The model displays two cross-kingdom-RNAi centric hypotheses to explain the observed reduction of the *metE* mRNA. Option (**I.**) involves the trafficking of exRNA including the identified miRNAs like let-7a-5p, which are potentially associated to human AGO proteins (hsAGO) and taken up by *A. fumigatus* to function in the fungus independently of its RNAi machinery. The other option (**II.**) surrounds an uptake of the human miRNAs and an incorporation of let-7a-5p (and other let-7 miRNAs) into PpdA/PpdB to silence *metE*. These hypotheses are not mutually exclusive. Model created in BioRender, Blango, M. (2027) https://BioRender.com/vjza2dy. (**B**) Total proteins were isolated from non-infected and conidia infected neutrophils as well as the correspondingly derived 20k and supernatant fraction. For the western blot, 5 µg of protein per lane was loaded and run on an SDS-polyacrylamide gel and further processed to detect all four human argonaute proteins, non-cleaving AGO1 and AGO4 (left panel) and cleaving AGO2 and AGO3 (right panel). Blue arrows indicate detected extracellular AGO-proteins. Detection of CANX served as a neutrophil loading control and verification, that the extracellular proteins are devoid of strictly intracellular content. (**C**) The *ppdA/B* double mutant (60) was incubated for 22 h and afterwards treated for additional 2 h with either 1X PBS (mock; dotted) or freshly isolated 20k fractions from non-infected (**-**) or infected neutrophils (**+**). After washing, the mycelium was harvested and subjected to RNA isolation, cDNA synthesis, and qPCR to quantify *metE*. (**D**) Control and mutant Let-7a-5p target site seed sequences in *metE* were incubated for 22 h and afterwards treated for additional 2 h as described above in (**C**). For (**C**) and (**D**), we removed two donors in each experiment that were non-responders under wild-type conditions to better assess the reversion of silencing. In both experiments, *metE* quantification was normalized to *cox5* mRNA levels (88). The statistical significance was calculated using a one-way ANOVA test followed by Šidák’s posttest controlling for multiple comparisons. A *p-*value 0.05 was considered significant.

In the second possible scenario let-7a-5p and other miRNAs hijack the *A. fumigatus* argonaute proteins PpdA/PpdB to guide them to the respective target mRNA (**Figure 3A, II.**). To test the fungal RNAi-independent option, we assessed the level of all human argonaute proteins in the extracellular fractions. Four AGO-proteins (AGO1-4) have been described for *Homo sapiens* of which only AGO2 and AGO3 retain their slicing capacity, while AGO1 and AGO4 are known to bear translation-blocking capability (60). Interrogation of total proteins from uninfected and infected neutrophils as well as the corresponding 20k and supernatant fractions by western blot resulted in different outcomes for all four proteins. While the blot for AGO2 confirmed earlier findings of no observable extracellular enrichment (**Figure 3B**; (37)), all other argonautes were detectable in the supernatant fraction at low levels. AGO1 displayed a constant extracellular abundance in the supernatant irrespective of the condition while the AGO4 amount decreased in the supernatant fraction during the *A. fumigatus* infection of neutrophils (**Figure 3B**). In contrast, AGO3 abundance was slightly increased in the infection-derived supernatant fraction (**Figure 3B**). Altogether, this confirmed an extracellular enrichment of three out of four argonaute proteins in the supernatant fraction.

While an uptake of the tested RNAi effectors to silence *metE* represents a valid option, the most efficient silencing was only detected with the infection-induced supernatant and 20k fractions, the latter not showing detectable AGO proteins. Accordingly, decreased *metE* mRNA levels might rely on the activity of the fungal RNAi machinery (**Figure 3A**, **I.**) and therefore be alleviated by the absence of the relevant proteins. To test this hypothesis we leveraged a Δ*ppdA/B* knockout strain devoid of any argonaute proteins, which was previously reported to have no hairpin-mediated RNA silencing activity (61). The double knockout strain was grown, treated, and processed the same way as the wild type and subjected to RT-qPCR quantifying *metE* transcript levels. We chose here to focus on the 20k fraction, as the Let-7 miRNAs were of highest relative abundance in this fraction. Challenging the wild-type and Δ*ppdA/B* hyphae with the 20k fraction derived during infection resulted in a similar decrease in comparison to the mock treatment (**Figure 3C**) similar to the earlier experiments (**Figure 2F**). As was already demonstrated (37), donor heterogeneity is observed in fungal responses to host extracellular fractions, and here we excluded two donors that did not exhibit silencing to better assess the reversion of silencing. Despite this donor heterogeneity, these data suggest that the decrease of *metE* mRNA levels does not rely only on fungal argonaute proteins and is instead likely conferred in a multifactorial manner, potentially involving both fungal and human RNAi machinery or other factors yet to be identified.

### Mutation of the let-7a-5p binding site in *metE* reverts silencing in a donor-dependent manner

We next wanted to confirm the validity of the predicted binding site of human let-7a-5p in *A. fumigatus metE* again using our strongest observed effect with infection-derived RNA of the 20k fraction. We created a strain harboring a chromosomal mutation of the predicted binding sites in the *metE* gene, disrupting the seed sequence of the miRNA:mRNA interaction (**Figure 3D**). We then repeated the extracellular fraction delivery experiments as before in **Figure 3C**. We observed a clear reversion of the *metE* silencing effect upon challenge with the infection-derived 20k fraction upon mutation of the *metE* binding sites in *A. fumigatus* (**Figure 3B**), despite observing no clear difference in *metE* expression between the control and mutated strains in mock treated samples from our RT-qPCR data. We again removed two non-responder donors in these assays. These data are consistent with a donor-dependent, cross-kingdom delivery of Let-7 family miRNA to silence gene expression in the dangerous human fungal pathogen *A. fumigatus*.

## DISCUSSION

Cross-kingdom RNAi has emerged as a widespread mechanism of gene expression manipulation between hosts and pathogens. Despite numerous examples in plants and insects (27, 29, 31, 32), this study represents, to our knowledge, the first example of cross-kingdom RNAi between a human immune cell and a filamentous fungal pathogen.

Using small RNA-seq, we were able to demonstrate a heterogeneous extracellular RNA landscape that is relatively impervious to infection status of host neutrophils, despite differences between the smaller supernatant fraction and the larger 20k fraction, particularly regarding tRNA-derived RNAs. Although no major differences were observed upon infection, we did notice that the larger 20k fraction was enriched for RNAs derived from the mitochondria, whereas the supernatant faction contained more nuclear-encoded tDRs and rDRs. The lack of differential RNA abundance suggests that additional factors, such as surface proteins, lipids, or even nucleic acids may contribute to targeting of these extracellular populations to fungal pathogens like *A. fumigatus*, consistent with the observations from [36] that host extracellular vesicles are altered upon the sensing of fungal dihydroxynaphthalene melanin. It would not be surprising that additional factors would be required for extracellular vesicle trafficking across the fungal cell wall. It has long been known that naked nucleic acids fail to enter the fungus in high levels, even when lipid encapsulated (60, 61). Such a targeting mechanism would be consistent with studies using RNA-loaded ambisomes (a lipid formulation of amphotericin B), which proved effective in facilitating delivery across the fungal cell wall, likely by supporting remodelling of the cell wall (62). Recent work in *Botrytis cinerea* showed that cell wall degrading enzymes are loaded onto fungal extracellular vesicles to facilitate entry into plant cells (63). Collectively, these data suggest that additional factors could be required to remodel the cell wall and facilitate trafficking.

Human neutrophils are known to undergo a range of responses to fungal pathogens, including degranulation, extracellular vesicle production, and even neutrophil extracellular trap formation (64, 65), resulting in a heterogeneous environment. It was not surprising then that our investigation of the extracellular RNA landscape of neutrophils revealed common RNA molecules like rRNA-derived RNAs, non-coding RNAs like the Y RNAs, and tRNA-derived RNAs. Interestingly, we noted differential loading of tRHs, e.g., valine, into different extracellular fractions, suggesting these tDRs might be useful as markers of these fractions. We took advantage of a sensitive stem-loop reverse transcription droplet digital PCR assay to confirm the production of tRNA-halves from several tRNAs in this system, noting that tDRs of even low abundance could be detected using this method. The ability to discriminate infected versus uninfected neutrophils using nuclear-encoded tDRs might then be an approach investigated for diagnostic purposes, as many studies are now underway with a variety of metabolic diseases and cancers to use tDRs as biofluid diagnostics (66). Future studies will be required to determine if biofluid exRNA composition could also prove useful in determining infection status. Likely, tDRs produced by neutrophils would not be the best source of diagnostic tDRs, as these cells are typically severely diminished in susceptible populations, but this approach nonetheless provides a proof-of-principle of tDR detection upon *A. fumigatus* infection.

In defining the extracellular RNA landscape, we noted enrichment of miR-223-3p in the supernatant fraction and Let-7 family miRNA in the 20k fraction of neutrophils. The abundance of miR-223-3p is likely linked to intercellular attenuation of the immune response in neighbouring cells (67), potentially during situations of neutrophil swarming (68). The Let-7 miRNAs were previously demonstrated to be antifungal in nature, silencing targets like *sec2* in insect-fungal pathogen and insect-bacterial interactions (32, 51). In the neutrophil-*A. fumigatus* interaction, we observed moderate silencing of *sec2* at the transcript level, consistent with a role for *sec2* in fungal biology (**Figure S9**; (52)). Future experiments should investigate a potential role in more robust post-transcriptional gene silencing. A stronger silencing effect was observed for *metE*, which had a higher sequence complementarity to the target sequence, which often leads to higher rates of endonucleolytic cleavage (14). Mutation of the *metE* seed sequence in *A. fumigatus* reverted the silencing effect (**Figure 3D**), consistent with a cross-kingdom RNAi phenotype. Future studies should more carefully investigate this molecular interaction between Let-7 miRNAs and *metE*.

To better understand the mechanism of silencing, we assessed the presence of human argonaute proteins in the 20k fraction but saw minimal protein at best. Surprisingly, deletion of the fungal argonautes did not revert the silencing effect (**Figure 3C**), suggesting that (i) low levels of human argonaute proteins are sufficient to mediate silencing, (ii) other fungal proteins contribute to a yet-to-be-described noncanonical silencing pathway as found in other fungi (69), or (iii) human and fungal proteins can interchangeably facilitate silencing, making the phenotype harder to assess. Another possibility is that an entirely alternative mechanism or indirect regulation leads to the decreased expression of *metE*. For example, we recently proposed that RNA:DNA:DNA triple helices may contribute to cross-kingdom gene expression (70), consistent with a known role for triplex formation by miRNAs (71). Future work will be required to formally test this hypothesis, but the presence of numerous lncRNAs and miRNAs in the neutrophil extracellular space is intriguing.

In conclusion, we provide support for the hypothesis that human neutrophils release extracellular RNAs that can serve as intercellular regulators, e.g., miR-223-3p, and function across kingdoms to regulate gene expression in the fungal pathogen *A. fumigatus*, e.g., let-7a-5p. Such a mechanism offers opportunities for potential intervention, for example in developing synthetic extracellular vesicles carrying necessary cargo to mediate gene silencing, even in the absence of immune cells like neutrophils in immunosuppressed patients (72). The data presented here also may serve as a starting point for improved liquid biopsy by revealing nuclear-encoded tDRs as a putative discriminatory exRNA. Ultimately, more effort in susceptible patient populations using human blood samples alongside refined methodology will be required to define the role of exRNA in the host response to pathogenic microbes.

## MATERIAL AND METHODS

### Ethics statement and neutrophil isolation from fresh venous blood

This study was conducted with the approval of the Institutional Review Board of the Jena University Hospital (approval number 2024-3324-Material) and in agreement with the Declaration of Helsinki. Anonymous healthy adult volunteers donated fresh venous blood for the performed experiments after granting informed and written consent. Polymorphonuclear granulocytes (PMNs; neutrophils) were isolated from fresh venous blood according to the method previously described in (37, 73) using PolymorphPrep solution (Progen) for the gradient centrifugation. Viability (≥90%) and the quantity of neutrophils was assessed with a Luna Cell Counter (Logos Biosystems).

### Cultures, growth conditions, and conidia opsonization

The *A. fumigatus* strains utilized in this study (**Table S4**) were grown on Malt Agar or *Aspergillus* minimal media (AMM, for transformations and droplet assays) at 37°C according to (60) for 5 days. Subsequently, after adding 10 mL sterile MiliQ water, conidia were harvested with a T-cell scraper and filtered with the help of a 30-μm cell strainer (MACS, Miltenyi Biotec GmbH) to remove remaining mycelium. The conidia suspension was centrifuged at 4000 rpm, 4°C for 5 min and the supernatant was removed while the pellet was taken up in 10 mL sterile MiliQ water. This washing step was repeated twice. The final 10 mL conidia suspension was quantified with a Roche Innovatis CASY Cell Counter and Analyzer System Modell TT.

Performing infection experiments with freshly isolated neutrophils and CEA10 conidia involved the opsonization of the latter according to (37) with few adjustments. Washed conidia were opsonized with 90% (vol/vol) 1X Phosphate Buffered Saline (PBS, Gibco^TM^ Thermo Fisher Scientific) and 10% (vol/vol) Normal Human Serum (NHS) for 1 h at 37°C, 500 rpm. Afterwards, the conidia were pelleted at 1600 x *g*, 4°C for 1 min and washed three times with 1X PBS. The final suspension was quantified with a Roche Innovatis CASY Cell Counter and Analyzer System Modell TT and stored on ice until further use.

Deletion of *sec2* (*AFUB_042620*) and *metE* (*AFUB_037120*) involved the generation of a knockout construct with the Gibson approach (New England Biolabs) following the manufacturer’s protocol. According to (60, 74), the gene deletion was performed by amplification of the selection marker from pSK485 (75) including about 1 kb of *sec2* or *metE* flanking sequences with the help of the primers given in **Table S5**. The resulting knockout cassette was employed for protoplast transformation and homologous recombination to delete the respective gene in *A. fumigatus* CEA17 Δ*akuB^KU80^* as described in (76), employing 1 M sorbitol-containing AMM agar plates that were supplemented with pyrithiamine (0.1 μg/ml) for positive clones selection. Where appropriate, excision of the selection cassette by the *xylP*-driven expression of β serine recombinase was performed by three rounds of transformant growth on 2% xylose-containing AMM. Subsequent growth control on pyrithiamine-containing AMM and a marker specific PCR were employed to validate the excision. Analytical DNA digestion and subsequent Southern blot were conducted to validate gene deletions or gene insertions in *A. fumigatus*. These approaches were performed according to (60, 74, 77).

A version of *metE* with targeted nucleotide exchanges to abolish let-7a-5p-miRNA binding without changing the amino acid sequence (from wt: 5’-AACGTTGACCACCTACTACCTC-3‘ to mut: 5‘-AACGTTGAC**A**ACCTA**T**TA**TT**TC-3‘ at position 876-882 of the CDS) was synthesised by Eurofins genomics. From this an insertion cassette was constructed consisting of a 1 kb flank downstream from the terminator of *metE*, *ptrA* resistance cassette, and the mutated sequence + a genomic targeting 1 kb flank. Essentially the same construct was created as a control but with a completely wild-type *metE* sequence. After transformation *metE*_mut was sequenced to ensure correct genetic insertion of the mutated sequence. Both the control and *metE*_mut were confirmed by Southern blot (**Figure S11B**).

### Neutrophil infection and exRNA isolation

Depending on the experimental set up, appropriate amounts of freshly isolated and quantified neutrophils (concentration ≥1 × 10^7^ cells/mL) were infected with opsonized wild-type CEA10 conidia (see above) with a MOI of 5 and incubated in Roswell Park Memorial Institute 1640 medium (RPMI1640, Gibco^TM^ Thermo Fisher Scientific) at 37°C, 5% CO_2_ (vol/vol) for 2 h. The mock treatment included the same amounts of cells being incubated under the same conditions without any fungal challenge. After the incubation the supernatant was removed and processed as described in [36] resulting in the 20k and supernatant fraction (**Figure 1A**). The different fractions from the respective conditions were subjected to RNA isolation with the miRNeasy Mini kit (QIAGEN) and quantified with the Qubit^TM^ RNA High Sensitivity (HS) assay kit and a Qubit flex fluorometer (Thermo Fisher Scientific) according to the manufacturer’s protocol. To ensure an efficient lysis, 100 µL glass beads (Ø 0.5 mm) were added to the samples and processed with a FastPrep-24™ homogenizer for 30 s at 4.0m/s with a 1 min break on ice, which was repeated two additional times. Depending on the experimental set up (e.g. tDRs quantification), Neutrophils were also subjected to the same RNA isolation kit and RNA was quantified with the Qubit^TM^ RNA Broad Range (BR) assay kit (Thermo Fisher Scientific).

### Small RNA identification with *miRge3.0*

Following library preparation with the Takara small RNA Library Preparation kit (SMARTer® smRNA-Seq Kit for Illumina®, Takara Bio Inc.) and sequencing with NovoGene, the fastq files were assessed for quality using FastQC (https://www.bioinformatics.babraham.ac.uk/projects/fastqc/). The fastq files were then processed using cutadapt (78) according to the specifications provided by Takara. The kit adds a polyA tail before the adapter, which requires removal of AAAAAAAAAA and everything that follows. 3 nucleotides were also removed from the 5′ end that are added in the template switching reaction to facilitate unbiased ligation. Ultimately this resulted in cleaner fastq files, as determined by fastqc. The supernatant and 20k experiments were performed separately as single-end 50 (SE50) and paired-end 150 (PE150) sequencing reactions, respectively. In both cases, single-stranded reads between 18 and 40 were kept for the final analysis.

The processed fastq files were aligned to the human genome (hg38) using *miRge3.0* (41) with options for -tRF and and -db miRGeneDB (79). The alignment to GRCh38 (hg38) was aided in annotation using GENCODE *Release 44* (80). The output was further processed using featureCounts (81) as part of the Rsubread package (82) to collect count data. These were processed by EdgeR to calculate TMM-normalized CPM values and determine differential expression using the 4 spontaneous and 4 infection-derived samples as the distinction (83), which allowed for consideration of fractional counts from multimapping reads. Downstream plots were created in R and Rstudio using packages ggplot2 (84), ggrepel (85), rtracklayer (82), and pheatmap (86) and information from **Table S1-2**. The supernatant and 20k fraction datasets were collected separately and treated separately throughout. ChatGPT version GPT-5.6 was used to support code development for creating plots downstream of RNA-seq read alignment. All resulting graphs and outputs were spot checked and cross-validated. Target prediction of identified human miRNAs against the *A. fumigatus* A1163 (ASM15014v1) cDNA library was performed using three tools, miRanda (S = 155, E = −20) (54), psRNATarget (default parameters of Schema V2, 2017 release) (55–57), and RNAhybrid (e = −20, gap = 2-8) (53).

### Stem loop-reverse transcription of tRNA-halves and droplet digital PCR

Quantification of candidate 5’ tRH employed the protocol from Tosar et al. (8, 9) to generate cDNAs with few adjustments. The stem-loop RT primers given in **Table S5** were pre-folded according to (87) and utilized in RT set ups with the Maxima H Minus First Strand cDNA-Synthesis-Kit (Thermo Fisher Scientific) and 5 ng total RNA input per sample with the following program: 16°C for 30 min, 50°C for 30 min, 85°C for 5 min. The generated cDNA was either diluted 1:3 to be tested in a standard PCR or 1:50 and 1:100 depending on the tested 5′ tRH for ddPCR quantification experiments. The PCR reaction was performed using the 2X MyRedTaq Master Mix (Meridian [Bioline]) with 0.4 µM per primer and the following program: **1.** 95°C, 3min; **2.** 95°C, 15s; **3.** 53°C-55°C, 15s; **4.** 72°C, 10s; **5.** Circle back to step 2., 24-34 cycles; **6.** 72°C, 5min; **7.** 8°C, ∞. Steps 3. and 5. were adjusted according to the utilized primer combination.

Droplet digital PCR was carried out using the QX200 ddPCR system (Bio-Rad) following the manufacturer’s recommendation. In brief, 20-µL reaction mixes were prepared containing 10 µL QX200 ddPCR Evagreen supermix (Bio-Rad) and the above-mentioned primer and DNA concentrations. After droplet generation with the QX200 droplet generator (Bio-Rad), PCR amplification was executed as followed: 95°C for 10 min; 40 cycles of 94°C for 30 s and 55°C for 1 min; 98°C for 10 min. Subsequently, the droplets were analyzed by the QX200 Droplet Reader (Bio-Rad) using the software Quanta Manager 1.2 Standard Edition. Only detections of more than 10,000 droplets per well were used for analysis.

### Hyphal treatment with extracellular content and RT-qPCR

Based on the sRNA-seq and let-7a-5p-target prediction data, *A. fumigatus* hyphal treatment experiments utilizing different strains were conducted. In general, 3 × 10^4^ conidia of the respective strains were seeded in a well of a 4-well dish with 3 mL of 2% glucose-adjusted RPMI 1640 for every tested condition (mock, 20k, and supernatant fraction from uninfected and infected conditions) and incubated for 22 h at 37°C, 5% CO_2_ (vol/vol). On the following day, extracellular fractions were isolated as described above and in (37) from 3 × 10^7^ neutrophils (uninfected and infected), whereas the supernatant fraction from both conditions were additionally concentrated with Amicon® Ultra-15 Centrifugal Filter Units (Merck Millipore). Prior to treatment, hyphae were washed once with 1X PBS after removing old media and replenished with 3 mL/condition of fresh glucose-adjusted RPMI 1640. Afterwards, the extracellular fraction isolates or 50 µL 1X PBS (mock treatment) were added to the corresponding hyphae, briefly mixed by careful shaking of the 4-well dish and incubated for additional 2 h at 37°C, 5% CO_2_ (vol/vol). The media of the treated hyphae was replaced with 2 mL 1X PBS and the fungal material was collected with a cell scraper to be transferred into a screw-cap tube prefilled with ∼100 µL of glass beads (Ø=0.5 mm). The hyphae were centrifuged at 12000 x *g*, 4°C for 5 min and the supernatant was discarded. This washing step was repeated two additional times. Finally, 700 µL of QIAzol (QIAGEN) were added to each sample and either directly subjected to RNA isolation or shock frozen with liquid nitrogen and stored at −70°C until further use.

The RNA isolation and quantification were conducted as given above with the mentioned adjustments. 300 ng of total RNA per sample were subjected to DNase I treatment (with RiboLock RNase inhibitor, Thermo Fisher Scientific) at 37°C, 1 h and after stopping the reaction by adding 1/10 of the reaction volume EDTA [50 mM] and incubation at 65°C, 5 min, samples were re-quantified with the Qubit^TM^ RNA BR assay kit (Thermo Fisher Scientific) and Qubit flex fluorometer. Subsequently, 100 ng of treated RNA isolates was subjected to cDNA synthesis using the Maxima H Minus First Strand cDNA-Synthesis-Kit (Thermo Fisher Scientific) according to the protocol for random hexamers and was quantified with the Qubit^TM^ ssDNA assay kit. For the qPCR experiments 2 ng cDNA per replicate were used with the Luna® Universal qPCR Master Mix (New England Biolabs) according to the manufacturer’s specifications. Experiments were performed with six biological replicates in technical triplicate and run on a CFX Duet Real-Time PCR System (Bio-Rad) using default settings for the threshold values (Ct). Calculation of the relative expression values was conducted with the 2^ΔΔ−CT^ method (88), normalizing the obtained data to the housekeeping gene *cox5* expression levels (60, 74).

### Western Blot

Total proteins were isolated from non-infected and conidia-infected neutrophils as well as the correspondingly derived 20k and supernatant fraction. For the western blot, 5 µg of protein per lane was loaded and run on an SDS-polyacrylamide gel. After wet transfer to PVDF membranes, membranes were blocked in blocking solution (1X PBS, 0.1% tween and 5% milk powder) at room temperature for 1 h, followed by incubation in respective primary antibody at 4°C overnight using the antibody dilution provided by the manufacturer (AGO1-4 sourced from rabbit and supplied by Cell signaling: AGO1, #9388; AGO2, #2897; AGO3, #5054; AGO4, #6913). ANXA antibodies were sourced from mouse, ANXA1 supplied by Revvity, #855301; ANXA7, #TA504925 and ANXA11, #TA500950 supplied by Origene. CANX sourced from rabbit and supplied by Cell Signaling, #2679). Membranes were washed with washing solution (1X PBS, 0.1% tween), followed by incubation with respective horseradish peroxidase (HRP)-conjugated secondary antibody (anti-rabbit [Thermo Fisher, #31460] or anti-mouse IgG [Thermo Fisher, #31430]) at room temperature for 2 h. Membranes were washed and signals were developed using Immobilon Western Chemiluminescence HRP Substrate (Sigma-Aldrich).

### Droplet Stress Assays

Potential growth phenotypes were analysed in response to specific challenges using droplet stress assays according to (74). The respective strains were spotted in serial dilutions of 10^2^ to 10^5^ on square petri dishes with 50 mL AMM, and if necessary, were supplemented with 1 mM diamide (Sigma-Aldrich), 0.5 µg/mL caspofungin (Biomol GmbH), 5 mM H_2_O_2_ (Carl ROTH), 0.3 mM 3-amino-1,2,4-triazole (3-AT) (Sigma-Aldrich), or 1 mM diamide + 0.5 µg/mL caspofungin. Incubation of the prepared plates was performed at 37°C with minimal disruption and documented with a Nikon D6500 AF-S NIKKOR 18–105 mm f / 3.5–5 ED VR camera after 48 h, 72 h, and/or 96 h.

### Statistical Analysis

Statistical analyses were performed using GraphPad Prism version 11.0.2 (92) or R version 4.5.0 (2025-04-11) in R Studio Version 2026.07.1+147 (2026.07.1+147) as described in detail in the specific methods sections or figure legends as appropriate.

## DATA AVAILABILITY

All raw and processed sequencing data generated in this study have been submitted to the NCBI Gene Expression Omnibus (GEO; https://www.ncbi.nlm.nih.gov/geo/) with accession numbers GSE348315 and GSE348316

## ACKNOWLEDGEMENTS

We would like to thank Kai Papenfort, Kathrin Fröhlich, Jorge Amich, and Amelia Barber for scientific discussions during this project. We would also like to thank Pamela Lehenberger, Nathalie Seiler, and Swatika Prabakar for technical support.

## FUNDING

The work presented here was generously supported by the Federal Ministry of Research, Technology and Space (BMFTR: https://www.bmftr.bund.de/), Germany, Project FKZ 01K12012 “RFIN – RNA-Biologie von Pilzinfektionen” and Project FKZ 01KI2515 “RFIN2.0– RNA-Biologie von Pilzinfektionen”. Additional funding support came from the Deutsche Forschungsgemeinschaft (DFG; German Research Foundation) under Germanýs Excellence Strategy – EXC 2051 – Project-ID 390713860. The funders had no role in the design, analysis, decision to publish, or preparation of the document.

## CONFLICT OF INTEREST

The authors declare no conflicts of interest.

## Notes

### Competing Interest Statement

The authors have declared no competing interest.

