## Supporting_Information for "Human neutrophils release extracellular RNA that modulates *Aspergillus fumigatus* gene expression across kingdoms"

**SUPPLEMENTAL TABLES**

**Table S1.** Extracellular RNAs detected in PMN-derived supernatant fraction as normalized read counts. See attached excel document.

**Table S2.** Extracellular RNAs detected in PMN-derived 20k fraction as normalized read counts. See attached excel document.

**Table S3.** miRNA target prediction output. See attached excel document.

**Table S4.** *Aspergillus fumigatus* strains used in this study. Given are the names of the strains, the genotype, and originating source of the strain.

| Strain | Genotype | Reference |
|---|---|---|
| **CEA10** | Wild Type | (1) |
| **CEA17 Δ*akuB*$^{KU80}$** | *akuB*$^{KU80}$*::pyrG*; PyrG+, Δ*akuB*$^{KU80}$ | (2) |
| ***Δsec2::ptrA*** | CEA17 Δ*akuB*$^{KU80}$; *AFUB_042620::six-xylP-ß-rec-trpC-T-ptrA-six* | This study |
| ***Δsec2::six*** | CEA17 Δ*akuB*$^{KU80}$; *AFUB_042620::six* | This study |
| ***ΔmetE::ptrA*** | CEA17 Δ*akuB*$^{KU80}$; *AFUB_037120::six-xylP-ß-rec-trpC-T-ptrA-six* | This study |
| ***ΔmetE::six*** | CEA17 Δ*akuB*$^{KU80}$; *AFUB_037120::six* | This study |
| ***ΔppdA/ppdB*** | CEA17 Δ*akuB*$^{KU80}$; *AFUB_038130::ptrA*; *AFUB_082250::ble* | (3) |
| **Mutated *metE*** | CEA17 Δ*akuB*$^{KU80}$; *AFUB_037120_mut::ptrA* | This study |
| **Control *metE*** | CEA17 Δ*akuB*$^{KU80}$; *AFUB_037120::ptrA* | This study |

**Table S5.** Oligonucleotides used in this study.

| Name | 5'-3' sequence | Target gene | Reference/ source |
|---|---|---|---|
| **Oligonucleotide primers for RT-qPCR** | | | |
| *metE*_o1 | TGCCGACGAGATGTTTGACA | *AFUB_037120*, mature mRNA | This study |
| *metE*_o2 | AGTGACGGCTACCAACGAAG | *AFUB_037120*, mature mRNA | This study |
| *AFUB_006820* _o1 | CAGACGTAGAGTCCATCGGC | *AFUB_006820*, mature mRNA | This study |
| *AFUB_006820* _o2 | CGTCGTCACCCTCATAGTCG | *AFUB_006820*, mature mRNA | This study |
| *AFUB_043410* _o1 | GCATGATCGCTCACGTTTCC | *AFUB_043410*, mature mRNA | This study |
| *AFUB_043410* _o2 | GGGGGTACATTTGGTGGAGG | *AFUB_043410*, mature mRNA | This study |
| *AFUB_081920* _o1 | CTTGCTGACCAACACTTGCC | *AFUB_081920*, mature mRNA | This study |
| *AFUB_081920* _o2 | CCGGGAGGTTGAGTTCTGAC | *AFUB_081920*, mature mRNA | This study |
| *sec2*_o1 | TTGCCAAGGACGATGTCGAA | *AFUB_042620*, mature mRNA | This study |
| *sec2*_o2 | GAGGAAAACTGGTCGAGGGG | *AFUB_042620*, mature mRNA | This study |
| **Oligonucleotide primers for cloning** | | | |

| | | | |
|---|---|---|---|
| *sec2_KO_o1* | GTCGACTCTAGAGGATCCCCTTAGCAGATCTTTCTCAGC | *AFUB_042620*, gene 5′ flank | This study |
| *sec2_KO_o2* | AGGCCTGAGTGGCCGGTACCGGCGGACTGGAGAGAGC | *AFUB_042620*, gene 5′ flank | This study |
| *sec2_KO_o3* | TGGCCATCTAGGCCGAGCTCAGACTCTTTTTCTGTTTTGC | *AFUB_042620*, gene 3′ flank | This study |
| *sec2_KO_o4* | GAATTCGAGCTCGGTACCCTGCGTATCGAACTGCTTG | *AFUB_042620*, gene 3′ flank | This study |
| *sec2_KO_o5* | GGGTACCGAGCTCGAATTC | Vector backbone | This study |
| *sec2_KO_o6* | GGGGATCCTCTAGAGTCGAC | Vector backbone | This study |
| *sec2_KO_o7* | GGTACCGGCCACTCAGGCCT | Resistance marker | This study |
| *sec2_KO_o8* | GAGCTCGGCCTAGATGGCCA | Resistance marker | This study |
| *sec2_KO_o9* | TTAGCAGATCTTTCTCAGCACTG | *AFUB_042620*, gene 5′ flank, for amplifying whole KO cassette | This study |
| *sec2_KO_o10* | TGCGTATCGAACTGCTTGATCCC | *AFUB_042620*, gene 3′ flank, for amplifying whole KO cassette | This study |
| *metE_KO_o1* | GTCGACTCTAGAGGATCCCCGCAACCTGCGTGAAATCGG | *AFUB_037120,* gene 5′ flank | This study |
| *metE_KO_o2* | AGGCCTGAGTGGCCGGTACCATGGCGAGTATACAGAG | *AFUB_037120,* gene 5′ flank | This study |
| *metE_KO_o3* | TGGCCATCTAGGCCGAGCTCAAGCCCACGACCTTCAT | *AFUB_037120,* gene 3′ flank | This study |

| | | | |
|---|---|---|---|
| *metE_KO_o4* | CGAATTCGAGCTCGGTACCCCTAGATGCT AAGCTCATTCGATTA | *AFUB_037120,* gene 3′ flank | This study |
| *metE_KO_o5* | CGAATGAGCTTAGCATCTAGGGGTACCG AGCTCGAATTCG | Vector backbone | This study |
| *metE_KO_o6* | ACCGATTTCACGCAGGTTGCGGGGATCC TCTAGAGTCGAC | Vector backbone | This study |
| *metE_KO_o7* | CTCTGTATACTCGCCATGGTACCGGCCAC TCAGGCCT | Resistance marker | This study |
| *metE_KO_o8* | CATATGAAGGTCGTGGGCTTGAGCTCGG CCTAGATGGCCA | Resistance marker | This study |
| *metE_mut_o1* | CCTGCAGCCCCTATAACGAATCATTCTTTC GCCGACATTG | *AFUB_037120,* gene 5′ flank together with mutagenized let-7a-5p binding site | This study |
| *metE_mut_o2* | TCCGCTTCCTCCTCTGTATACTCGCCATGG CT | *AFUB_037120,* gene 5′ flank together with mutagenized let-7a-5p binding site | This study |
| *metE_mut_o3* | ATGGAAAAACAATAGCTGGCTGGACTCG AG | *AFUB_037120,* gene 3′ flank | This study |
| *metE_mut_o4* | AATCCACATGTCAACCGGGAGAAGGGTG | *AFUB_037120,* gene 3′ flank | This study |
| *metE_mut_o5* | TATACAGAGGAGGAAGCGGAAGAGCGC | Vector backbone | This study |
| *metE_mut_o6* | GCCAGCTATTGTTTTTCCATAGGCTCCGC CC | Vector backbone | This study |

| | | | |
|---|---|---|---|
| *metE_mut_o7* | TCCCGGTTGACATGTGGATTACGAGCTAACGAGT | Resistance marker | This study |
| *metE_mut_o8* | TTCGTTATAGGGGCTGCAGGAATTCGATGG | Resistance marker | This study |
| *metE_ctrl_o1* | TATACAGAGGAGGAAGCGGAAGAGCGC | metE_mut plasmid without 5′ flank | This study |
| *metE_ctrl_o2* | TTCGTTATAGGGGCTGCAGGAATTCGATGG | metE_mut plasmid without 5′ flank | This study |
| *metE_ctrl_o3* | CCTGCAGCCCCTATAACGAATCATTCTTTCGCCGACATTG | *AFUB_037120,* gene 5′ flank, together with natural let-7a-5p binding site | This study |
| *metE_ctrl_o4* | TCCGCTTCCTCCTCTGTATACTCGCCATGGCT | *AFUB_037120,* gene 5′ flank, together with natural let-7a-5p binding site | This study |
| **Oligonucleotide primers – Stem-loop RT-PCR** | | | |
| Universal reverse primer | GTGCAGGGTCCGAGGT | Stem-loop RT-primer | (4) |
| hs-tRNA$^{Asp}_{GTC}$ 5´half_Fw | ggTCCTCGTTAGTATAGTGGTGAG | hs-tRNA$^{Asp}_{GTC}$ 5´half | This study |
| hs-tRNA$^{Val}_{CAC}$ 5´half_Fw | ggGTTTCCGTAGTGTAGTGGT | hs-tRNA$^{Val}_{CAC}$ 5´half | This study |
| hs-tRNA$^{Glu}_{TTC}$ 5´half_Fw | TCCCACATGGTCTAGCGGT | hs-tRNA$^{Glu}_{TTC}$ | This study |

| | | | |
|---|---|---|---|
| | | 5´half | |
| hs-tRNA$^{Asp}_{GTC}$ 5´half_SL | GTCGTATCCAGTGCAGGGTCCGAGGTAT TCGCACTGGATACGACcaggcg | hs-tRNA$^{Asp}_{GTC}$ 5´half | This study |
| hs-tRNA$^{Val}_{CAC}$ 5´half_SL | GTCGTATCCAGTGCAGGGTCCGAGGTAT TCGCACTGGATACGACaggcga | hs-tRNA$^{Val}_{CAC}$ 5´half | This study |
| hs-tRNA$^{Glu}_{TTC}$ 5´half_SL | GTCGTATCCAGTGCAGGGTCCGAGGTAT TCGCACTGGATACGACaaaacc | hs-tRNA$^{Glu}_{TTC}$ 5´half | This study |

### SUPPLEMENTAL FIGURES

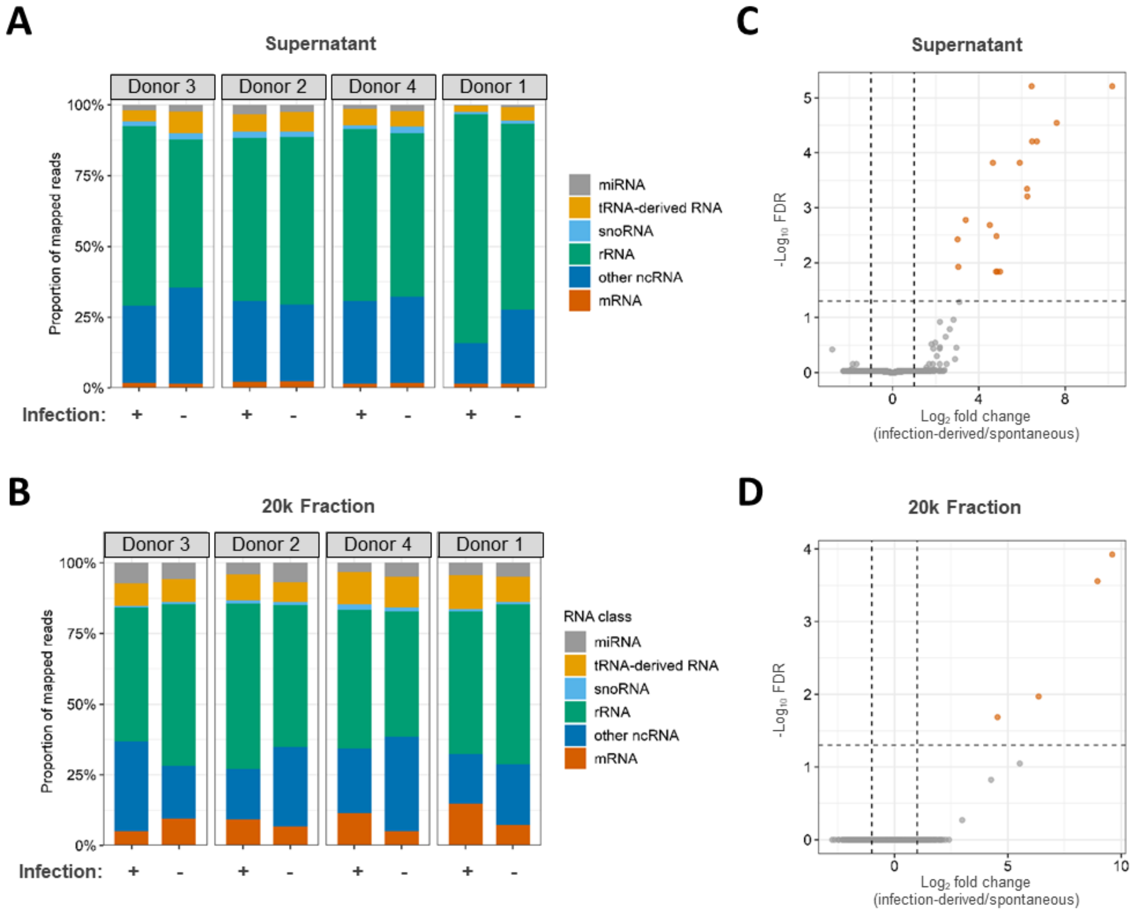


Figure S1. The extracellular RNA-ome is comprised with a variety of RNA species. Small RNA-seq combined with an *miRge3.0* analysis uncovered the composition of RNA species found in the neutrophils extracellular space during non-stressed and *A. fumigatus* stressed conditions among both the (**A**) supernatant and (**B**) 20k fractions for four human donors. Different colors indicate identified small RNA species. (**C-D**) Volcano plots of differentially expressed total RNAs using all RNAs found in the analysis. A gene had to exhibit greater than 2-fold change and a Benjamini-Hochberg-adjusted *p*-value less than 0.05 to be considered significant.

**A**

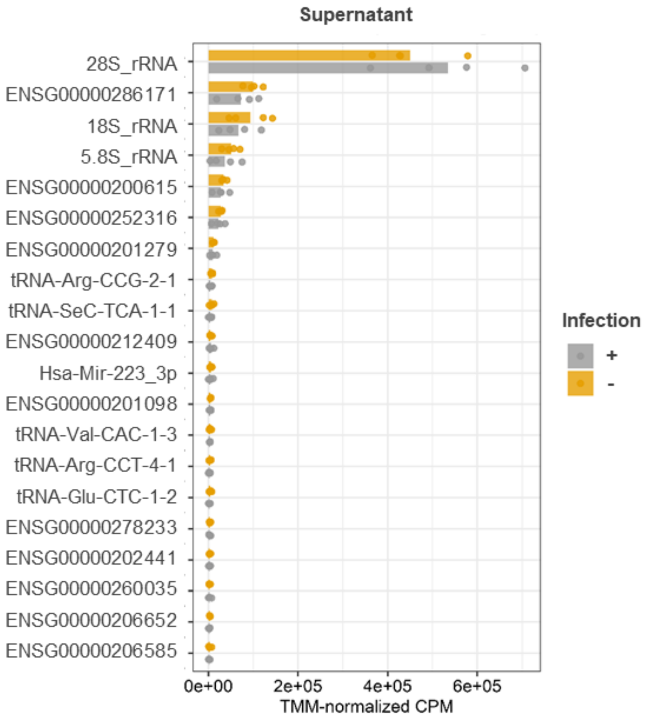


**B**

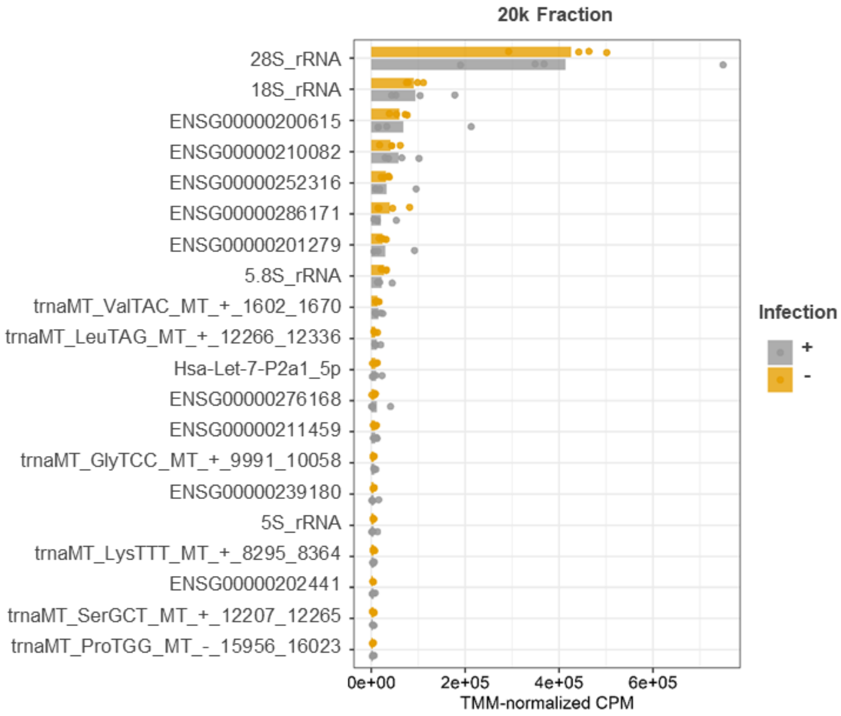


Figure S2. The extracellular RNAome contains RNAs derived from numerous sources. (**A-B**) The top 20 most abundant RNAs found in the supernatant (**A**) and 20k fraction (**B**) using *miRge3.0* software tool.

**A**

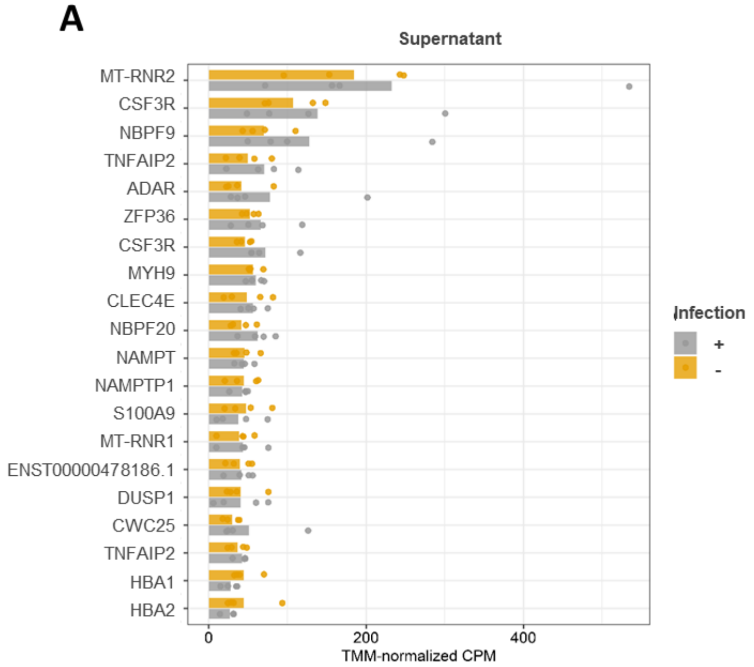


**B**

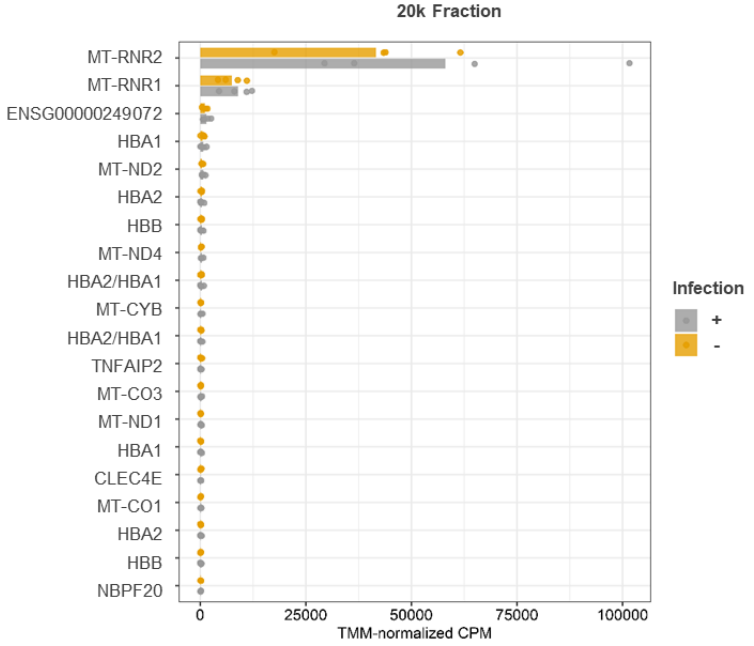


Figure S3. mRNA-derived small RNAs are differentially abundant in the supernatant fraction (**A-B**) Top 20 most abundant mRNA-derived transcripts for supernatant (**A**) and 20k fractions (**B**), respectively.

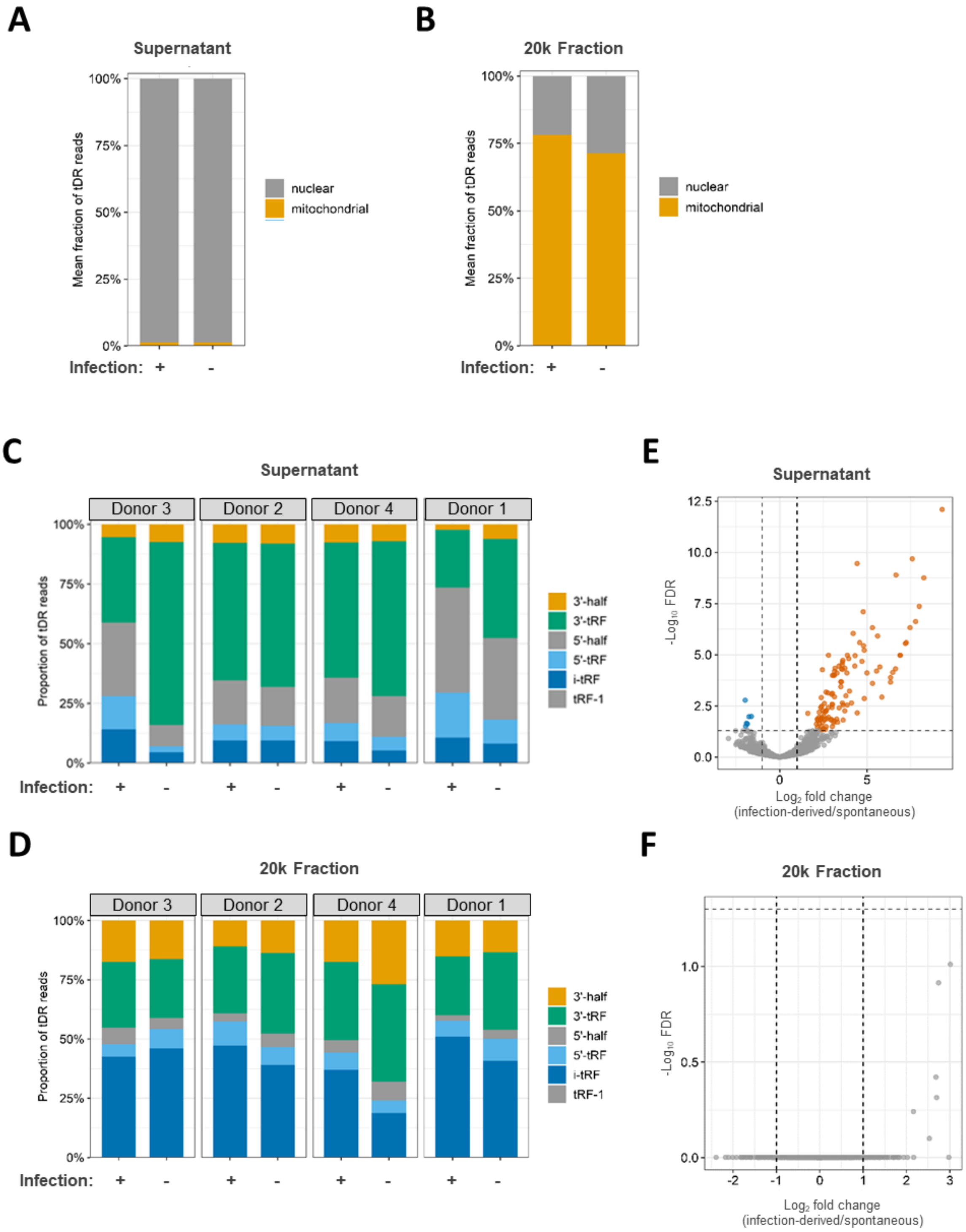


Figure S4. Supernatant and 20k fractions harbor different compositions of tRNA-derived RNAs. (**A-B**) Source of tDRs in supernatant versus 20k fractions using *miRge3.0* analysis. (**C-D**) Biotype composition of tDRs from four human donors and associated volcano plots (**E-F**). A tDR had to exhibit greater than 2-fold change and a Benjamini-Hochberg-adjusted *p*-value less than 0.05 to be considered significant.

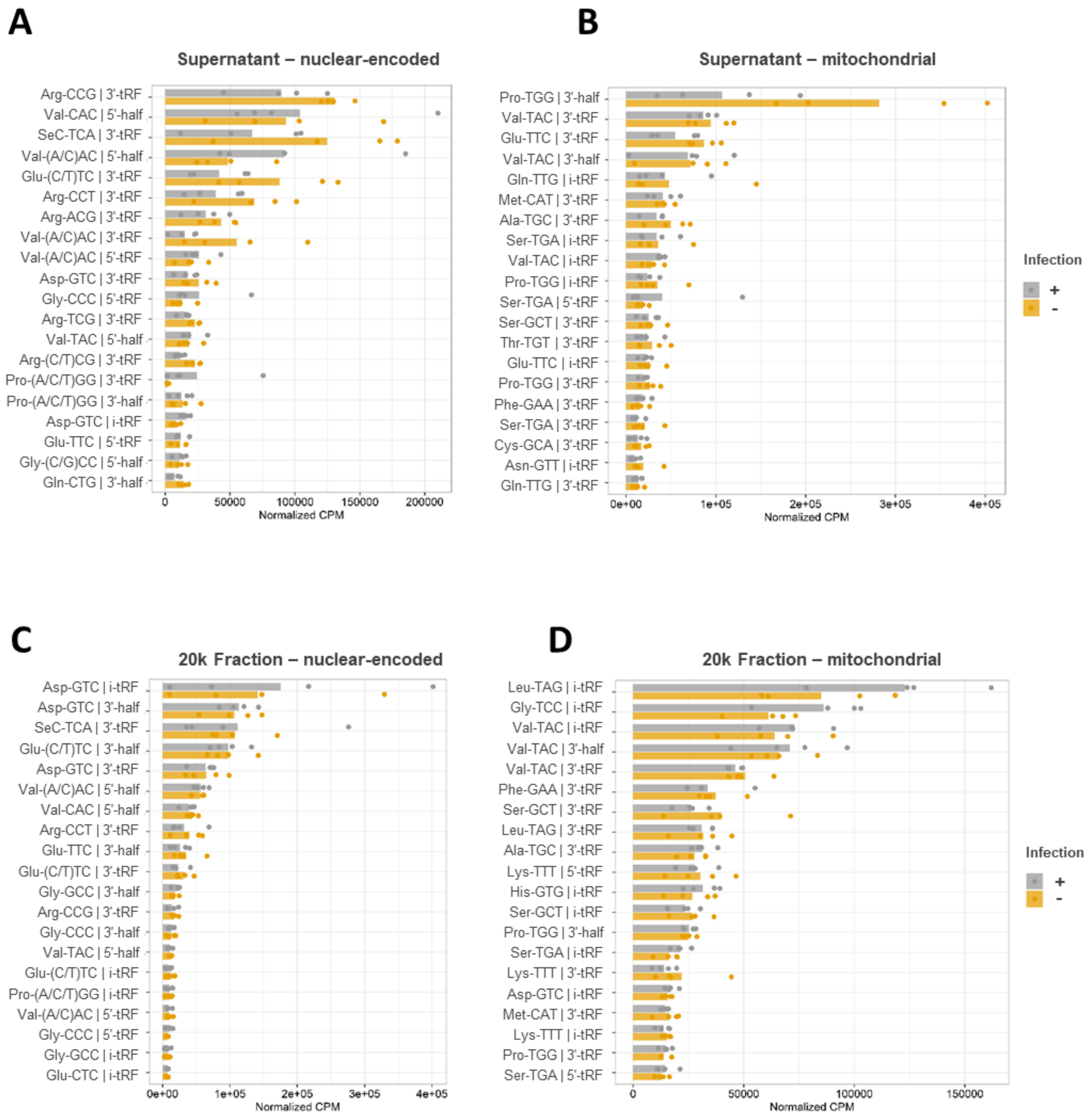


Figure S5. (**A-B**) Top 20 tDRs by isoacceptor class for nuclear-encoded tDRs and mitochondria-encoded tDRs of the supernatant fraction, respectively. (**C-D**) Top 20 tDRs by isoacceptor class for nuclear-encoded tDRs and mitochondria-encoded tDRs of the 20k fraction, respectively.

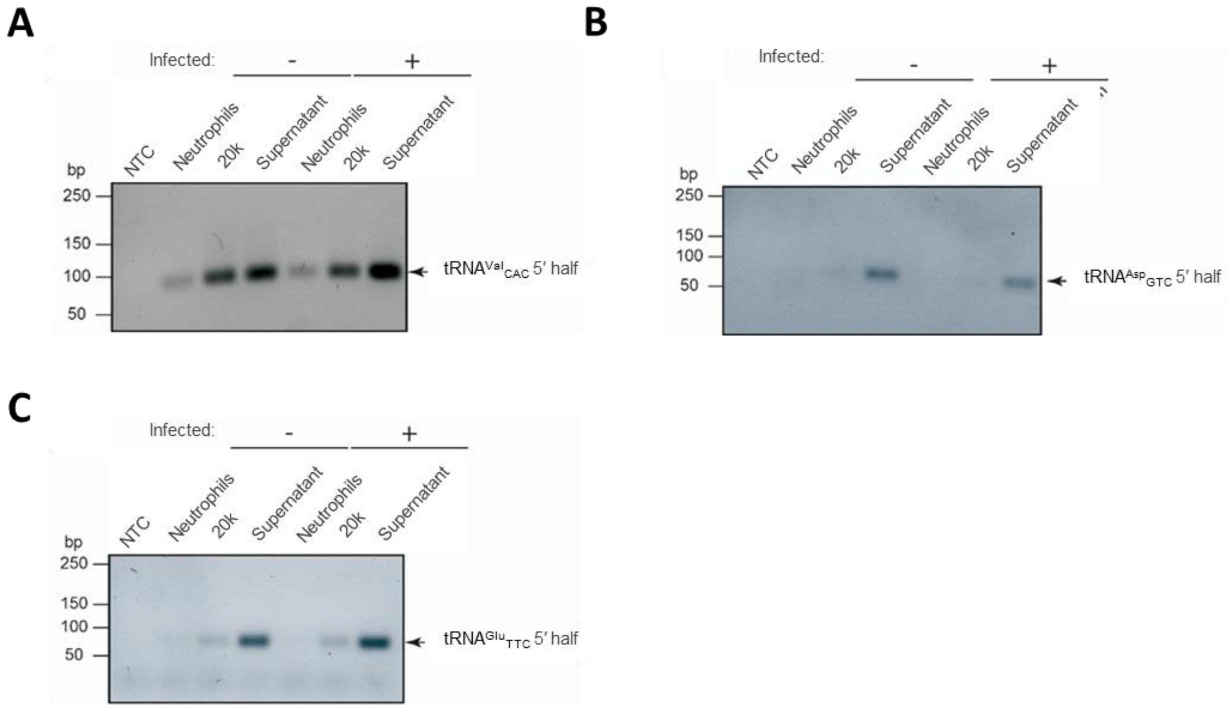


Figure S6. Application of a stem loop-primer based reverse transcription approach allows for the quantification of specific candidate tRFs. Using the protocol from Tosar *et al*. (4, 5), total RNA from PMNs and extracellular samples from both conditions were subjected to the SL-RT approach testing for three candidate 5’tRHs ($tRNA^{Val}_{CAC}$, $tRNA^{Asp}_{GTC}$, and $tRNA^{Glu}_{TTC}$). The generated cDNA was tested in a standard PCR with the respective primer combinations given in **Table S1** before measuring the samples with a ddPCR platform.

A

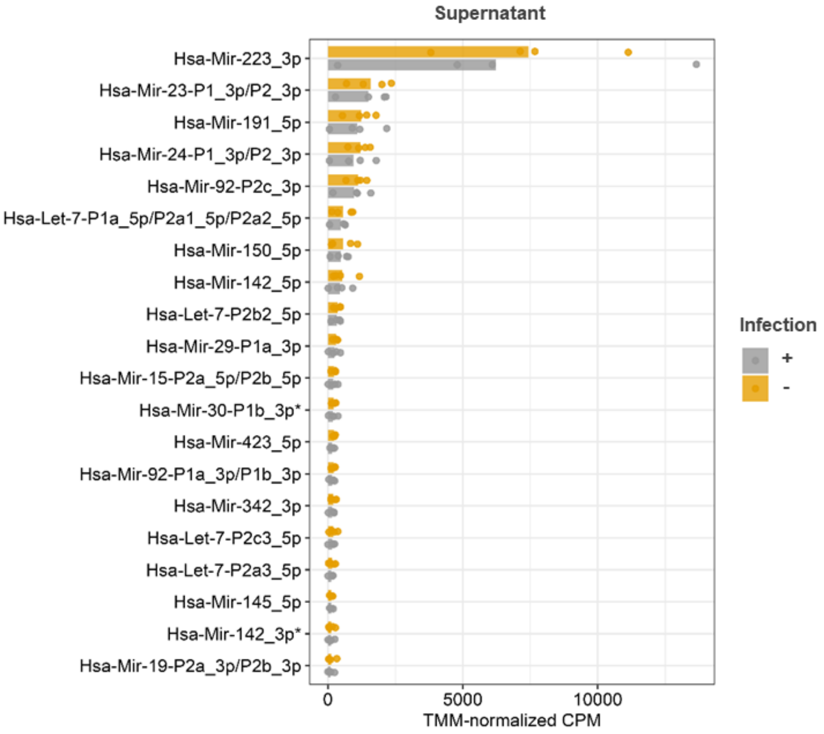


B

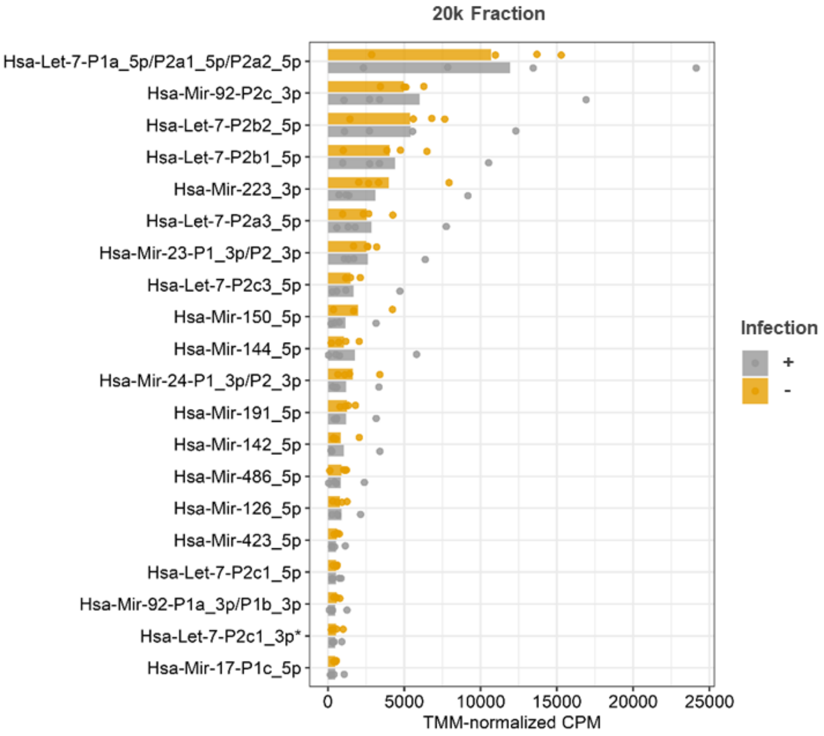


Figure S7. (**A**) Top 20 extracellular miRNA of the supernatant fraction for infected-derived and uninfected PMNs using *miRge3.0* software. (**B**) Top 20 extracellular miRNA of the 20k fraction for infected-derived and uninfected PMNs.

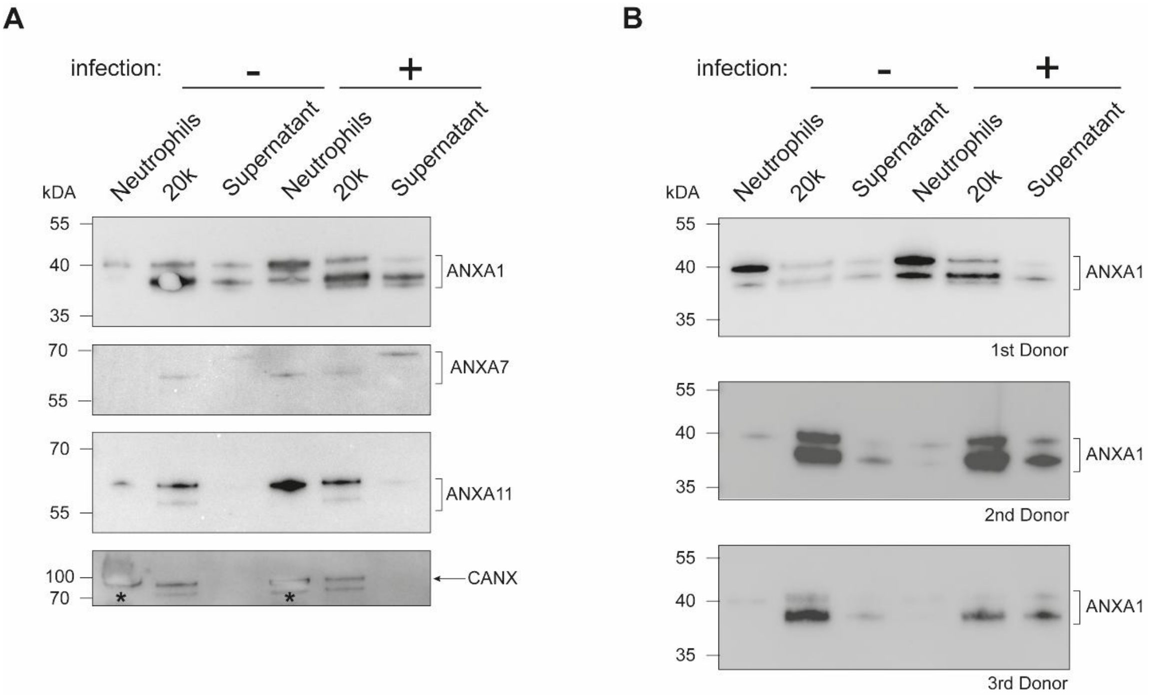


Figure S8. Extracellular enrichment of ANXA1, ANXA7, and ANXA11 is observed for *A. fumigatus* infected neutrophils. Total proteins were isolated from non-challenged and *A. fumigatus* challenged neutrophils together with the corresponding 20k and supernatant fraction. The samples were subjected to SDS-PAGE and western blot utilizing 5µg protein per lane to detect ANXA1, ANXA7, and ANXA11 (**A**) or ANXA1 in additional donor samples (**B**). CANX served as control to differentiate between intracellular/extracellular protein content. The asterisk indicates oversaturated signal intensity resulting in a blurry band.

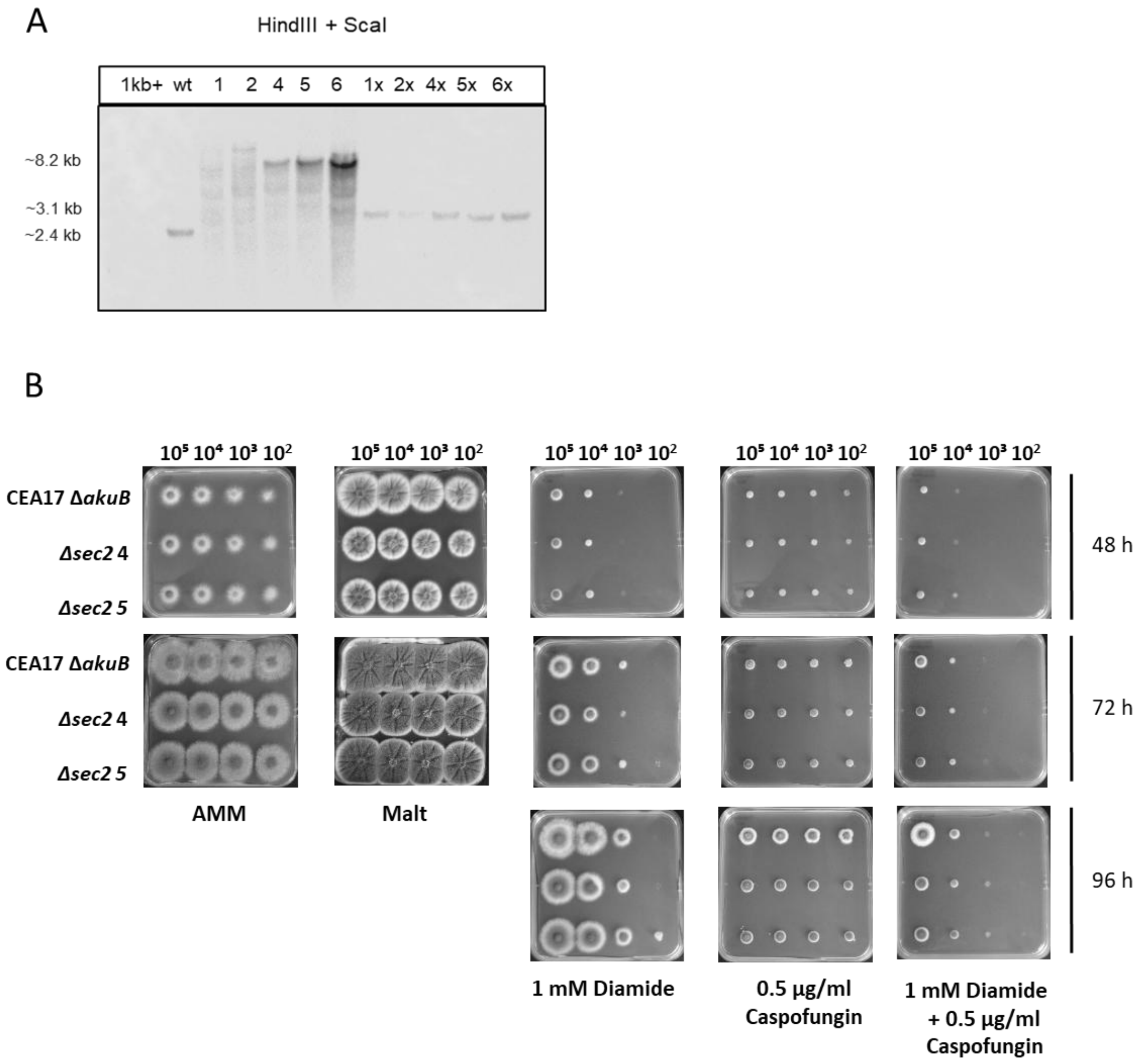


Figure S9. Deletion of *A. fumigatus sec2* resulted in decreased growth on malt agar while increasing susceptibility against caspofungin. (**A**) Southern blot confirming digestion pattern of the *sec2* knockout using *Hind*III and *Sca*I with DIG-labelled flanking region as probe. Successful insertion of the knockout cassette is shown by a digestion product of 8282 bp (with marker) and 3088 bp (without marker), while wild-type strain gives a digestion product of 2408 bp. (**B**) The indicated strains were spotted on AMM agar plates containing either 1 mM diamide and/or 0.5 µg/mL caspofungin. Additionally, droplet assays on malt agar and AMM agar of the wild type (WT) and two clones of Δ*sec2::ptrA* and Δ*sec2::six*, the selection marker being rescued for the latter one, served as control. Documentation was conducted after 48 h and 72 h of incubation at 37°C. The depicted assays are representative images of three biological replicates.

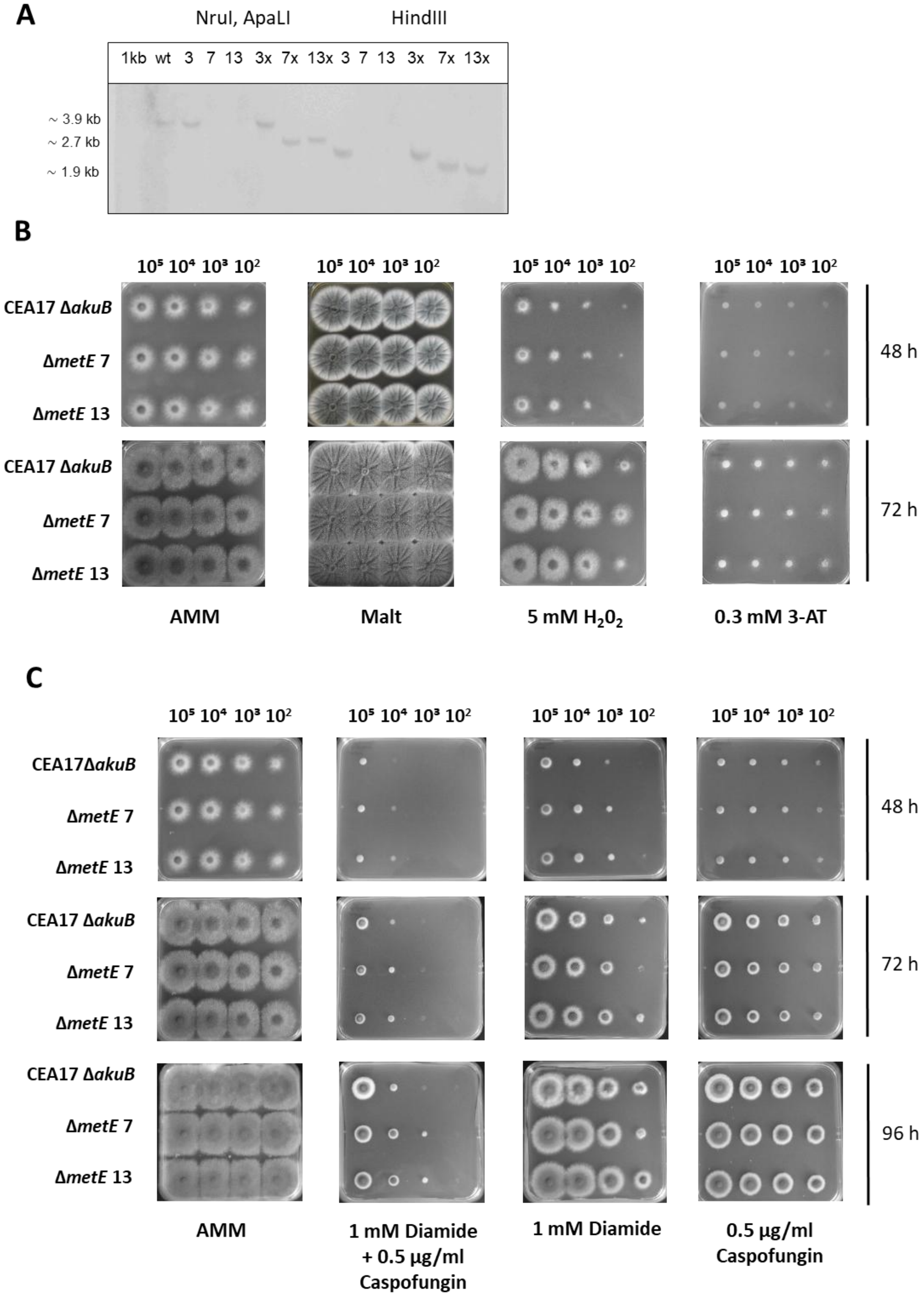


Figure S10. Deletion of *A. fumigatus metE* resulted in decreased growth under combination stress. (**A**) Southern blot confirming digestion pattern of the *metE* knockout using *Nru*I/*Apa*LI and *Hind*III with DIG-labelled flanking region as probe. Successful insertion of the knockout cassette is shown by a *Nru*I/*Apa*LI digest, yielding a digestion product of 7926 bp (with marker) and 2732 bp (without marker), while wild type strain gives a digestion product of 3929 bp. Successful distinction of marker containing and marker free strains is further verified by *Hind*III digest, yielding a digestion product of 7935 bp (with marker) and 1941 bp (without marker). The indicated strains were spotted on AMM agar plates containing either (**B**) 5 mM $H_2O_2$ or 3-amino-1,2,4-triazole (3-AT) or (**C**) 1 mM diamide and/or 0.5 µg/mL caspofungin. Additionally, droplet assays on malt agar and AMM agar of the wild type (WT) and two clones of Δ*sec2::ptrA* and Δ*sec2::six*, the selection marker being rescued for the latter one, served as control. Documentation was conducted after 48 h and 72 h of incubation at 37°C. The depicted assays are representative images of three biological replicates.

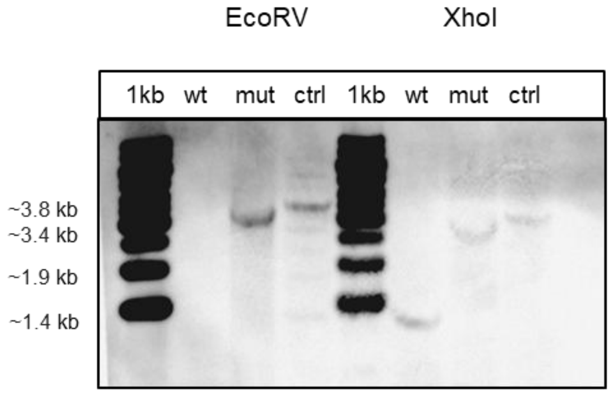


Figure S11. Southern blot confirmation of *metE* mutant and control strains using *Eco*RV and *Xho*I enzymes for digestion and a DIG-labelled flanking region as probe. Successful insertion of the mutagenized/control cassette is shown by *Eco*RV digest, yielding a digestion product of 3864 bp, compared to the wild-type strain which yields a product of 1946 bp. The insertion of the mutagenized/control cassette is further confirmed by a *Xho*I digest, yielding a digestion product of 3373 bp, compared to the wild type strain which yields a product of 1455 bp.

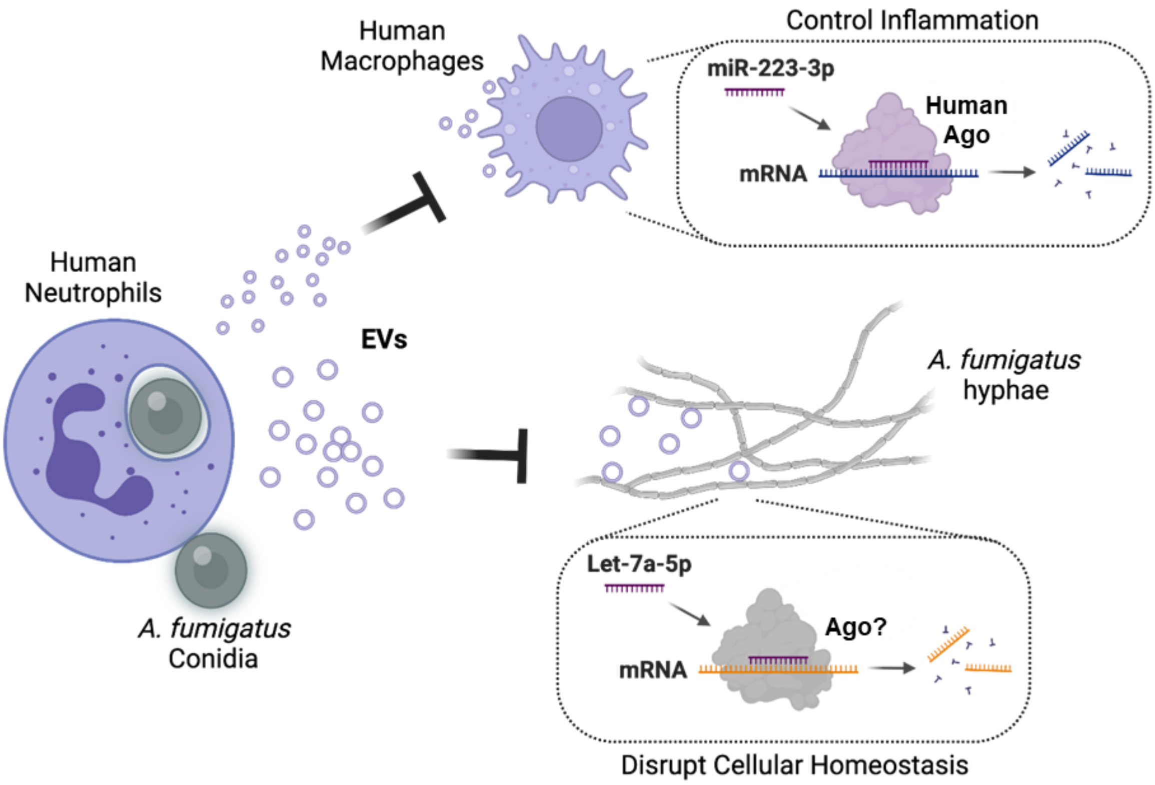


Figure S12. Working model of PMN-derived extracellular RNA trafficking in response to *A. fumigatus* infection. Model created in BioRender, Blango, M. (2027) https://BioRender.com/0tt7oup.
